# Effects of prenatal maternal immune activation and exposure to circadian disruption during adolescence: exploring the two-hit model of neurodevelopmental disorders

**DOI:** 10.1101/2024.02.25.580567

**Authors:** Tara C. Delorme, Danusa M. Arcego, Danae Penichet, Nicholas O’Toole, Nikki Huebener, Patrícia P. Silveira, Lalit K. Srivastava, Nicolas Cermakian

**Author notes:** Co-corresponding authors **Correspondence requests to:** Dr. Nicolas Cermakian, 6875 Boulevard LaSalle, Montréal, QC H4H 1R3, Canada; Dr. Lalit Srivastava, 6875 Boulevard LaSalle, Montréal, QC H4H 1R3, Canada.

## Abstract

**Background:** Around 80% of individuals with neurodevelopmental disorders (NDDs) such as schizophrenia and autism spectrum disorders experience disruptions in sleep/circadian rhythms. We explored whether prenatal infection, an established risk factor for NDDs, and environmental circadian disruption synergistically induced sex-specific deficits in mice.

**Methods:** A maternal immune activation (MIA) protocol was used by injecting pregnant mice (at E9.5) with a viral mimic poly IC or saline. Then, juvenile/adolescent offspring (3-7 weeks old) were subjected to either standard lighting (12:12LD) or constant light (LL).

**Results:** We found interactions of the two factors on behaviors related to cognition, anxiety, and sociability. Also, poly IC exposure led to a more activated profile of hippocampal microglia in males only, while LL diminished these effects. Using RNA sequencing in the dorsal hippocampus, we found that poly IC exposure led to many differentially expressed genes in males (but not females), and fewer differentially expressed genes were observed after LL exposure. Using the WGCNA analysis, we found several significant gene modules positively associated with poly IC (in comparison to saline exposure) and LL (in comparison to LD exposure) in males, and less so in females. Interestingly, many of the identified hub bottleneck genes were homologous to human genes associated with both sleep/circadian rhythms and neurodevelopmental disorders as identified by GWA studies.

**Conclusions:** Our work demonstrates that in a mouse model of prenatal infection, disruptions in circadian rhythms induced by LL play a role in modulating the effects of MIA at behavioral, cellular, and molecular levels.

## Introduction

Schizophrenia (SCZ) and autism spectrum disorders (ASDs) are neurodevelopmental disorders, each affecting approximately 1% of the global population(1, 2). Neurodevelopmental disorders result from complex interactions between genetic and environmental risk factors, which are thought to disturb normal brain development, ultimately leading to the clinical occurrence of these disorders(3–5). Current research increasingly focuses on the combined impact of multiple risk factors for neurodevelopmental disorders, rather than examining them in isolation(6).

A significant risk factor to consider is prenatal infection, which is linked to an increased risk of neurodevelopmental disorders in offspring(7, 8). Specifically, prenatal infection during the first trimester elevates the likelihood of offspring developing SCZ (9) and ASDs(10). Prenatal infection is thought to act as a primer for disease, potentially interacting with other risk factors to trigger the full array of symptoms(11). A maternal immune activation (MIA) protocol in animals, using polyinosinic:polycytidylic acid (poly IC), a viral mimetic, can be used to model prenatal infection(12).

In utero MIA leads to adult mouse offspring displaying behavioral deficits related to neurodevelopmental disorders, such as reduced sociability(13, 14), and cellular changes, in particular, altered microglial properties(15). Prenatal immune activation may affect microglia and alter their function in the developing brain, which can directly contribute to behavioral, cellular, and molecular deficits observed in MIA models(16, 17). MIA may also prime microglia to respond more or less to immune challenges experienced later in life (18, 19).

A less frequently discussed factor in the context of neurodevelopmental disorder is the disruption of circadian rhythms. Circadian rhythms are daily cycles lasting about 24 hours in behavior (e.g., sleep) and physiology (e.g., certain hormones and genes). These rhythms are generated through clock mechanisms found in most mammalian cells(20). To maintain synchrony with the environment, these endogenously generated rhythms require daily “resetting” by cyclic environmental cues, the strongest being light exposure(21). Disruption of the circadian timing system, often caused by inappropriate light exposure, can have adverse effects on both mental and physical health, especially when experienced chronically(22, 23). Notably, ∼80% of individuals with SCZ and ASDs exhibit disruptions in sleep, rest/activity and molecular rhythms, daily hormone rhythms and circadian clock gene expression(24–27). Circadian disturbances were reported in animal models based on genetic risk factors for SCZ (28) and ASDs (29) and in poly IC-exposed mice(30). Interestingly, in many individuals with SCZ, circadian disruptions precede the onset of psychosis(31, 32), suggesting that circadian disruption may be a risk factor for neurodevelopmental disorders. This hypothesis is supported by some mouse models for SCZ, where SCZ-related behaviors worsened after altered light exposure(33–35).

The current study explored if the combination of MIA as a first risk factor in-utero and exposure to circadian disruption as the second risk factor in adolescence would synergistically lead to behavioral deficits, alterations in microglial morphology and density and changes in gene expression in offspring. Adolescence was hypothesized to be the age most susceptible to the effects of circadian disruption because adolescence is already characterized by prevalent circadian disruption, including a shift towards later sleep/waking times and delayed onset of biological rhythms(36), which often leads to weekday sleep deprivation and increased weekend sleep(37). Additionally, adolescents frequently experience nighttime light exposure from electronic devices used for reading, socializing and entertainment(38, 39). Due to significant sex differences observed in SCZ(40), ASD (41) and in the MIA animal model(42), we incorporated male and female mice in our experiments. We found an interaction of MIA and circadian disruption at the behavioural, cellular (microglia) and molecular (transcriptome) levels.

## Materials and Methods

### Animals

C57BL/6J mice (Jackson Laboratory; product number: 000664) were used in a mating protocol described in the next section. MIA offspring and controls were housed in light-proof, ventilated cabinets with a light intensity of 150-200 lux (cool white LED lighting) controlled by an external timer (Actimetrics, Wilmette, IL, USA). For the three-chamber social interaction test, male and female C57BL/6 mice to be used as “strangers” were obtained from Charles River Laboratories (Saint-Constant, QC, Canada) and matched for age and sex with the test mice. Animal use adhered to the Canadian Council of Animal Care guidelines and was approved by the Douglas/McGill University Animal Care Committee.

### Maternal immune activation (MIA) protocol

The MIA protocol was done as described(30). In short, 8-week-old male and female mice were bred, and on embryonic day 9.5, pregnant dams were intraperitoneally injected with poly IC (5 mg/kg; lot: 059M4167V; Sigma-Aldrich, St. Louis, MO, USA) or sterile saline. Dams were left undisturbed to deliver their litters. Litters of poly IC- and saline-exposed dams did not differ in number of pups (Supplementary Fig. 1A, D, G).

In the first experiment (measuring various behaviors), 29 female breeders were mated with 15 male breeders: 3 dams were not pregnant, 11 out of 14 poly IC-injected dams gave birth (average litter size: 6.44, total male offspring used= 35, total female offspring used = 22) and 10 out of 12 saline injected dams gave birth (average litter size: 6.55, total male offspring used = 33, total female offspring used = 25). For the second experiment (assessing microglial morphology and density), 18 female breeders were mated with 9 male breeders: 2 dams were not pregnant, 6 out of 9 poly IC-injected dams gave birth (average litter size: 7.25, total male offspring used = 29, total female offspring used = 28) and 6 out of 7 saline-injected dams gave birth (average litter size: 6.4, total male offspring used = 13, total female offspring used = 11). For the third experiment (measuring gene expression using RNA sequencing), 20 female breeders were mated with 10 male breeders: 5 dams were not pregnant, 4 out of 8 poly IC-injected dams gave birth (average litter size: 7.5, total male offspring used = 14, total female offspring used = 16) and 7 out of 7 saline-injected dams gave birth (average litter size: 6.9, total male offspring used = 27, total female offspring used = 21).

Cytokines and chemokines were measured in a separate cohort of pregnant dams to validate that the lot of poly IC induced an immune response. Briefly, trunk blood was collected from pregnant dams (n=3/treatment) 3 hours after poly IC or saline injection. The Mouse Cytokine Array Proinflammatory Focused 10-plex (MDF10, Eve Technologies, Calgary, AB, Canada) was used for the following pro/anti-inflammatory biomarkers: IL-6, IL-2, IL-1β, IL-10, TNFα, MCP-1, IL-4, IL-12p70, IFNγ and GM-CSF (Supplementary Fig. 2A-I). Samples that were below the detectable range were designated as 0 pg/mL, and data for IFNγ are not reported because all samples were below the minimum detectable concentration.

### Experimental timeline

The experimental protocol is depicted in Fig. 1. Poly IC- and saline-exposed male and female offspring were aged to 3 weeks old, weaned and placed into light-proof, ventilated cabinets. Mice were given 2 days to habituate, before being subjected to 4 weeks of either a standard 12 h light: 12 h dark cycle (12:12LD) or constant light (LL) (i.e., from 3 weeks to 7 weeks old, equivalent to the juvenile/adolescence period). At 7 weeks, separate groups of mice were used for each of the following three experiments. The first cohort underwent behavioral testing under dim light conditions, where the lighting intensity at the center of the testing chambers was approximately 10 lux (except for Barnes maze). In the second cohort, brains were harvested for immunohistochemistry to analyze microglia in the prefrontal cortex, dentate gyrus, and the CA1 region. In the third cohort, the hippocampus was microdissected, and bulk RNA sequencing was performed.

**Fig. 1.**
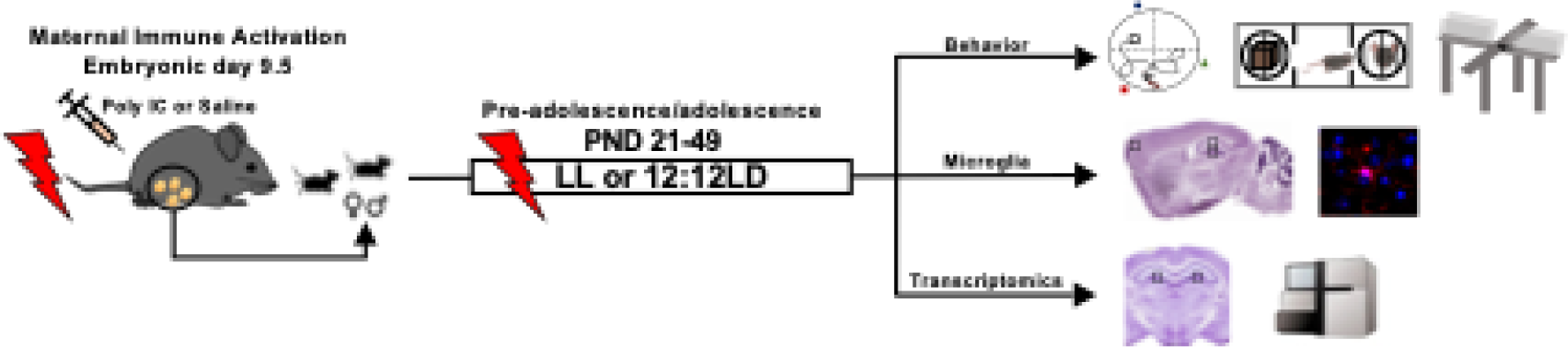
Experimental timeline. Pregnant dams were intraperitoneally injected with poly IC or saline at E9.5. Poly IC- and saline-exposed offspring were aged to 3 weeks old, then subjected to 4 weeks of either standard lighting (12:12LD) or constant light (LL). At 7 weeks, separate groups of mice were used for each of the three experiments: behavioral testing under dim light conditions; harvesting of brains for immunohistochemistry to analyze microglia in the prefrontal cortex, dentate gyrus, and CA1 region; microdissection of hippocampus, used for RNA sequencing.

### Behavioral outcomes

Behavioral tests were conducted as described previously(35). For each test, 16-18 males and 10-14 females were used per treatment (MIA or saline), and per lighting condition (LD or LL). Due to practical constraints, a subset of mice was randomly chosen for the social interaction test (n=8/group) and the Barnes maze (n=9-11/group). Tests were performed between 1 hour and 3 hours after lights on [Zeitgeber Time (ZT) 1-3]. In the LL condition, given that mice free ran (i.e., were not entrained to external time cues), passive infrared sensors were used to measure locomotion. Tests under LL were conducted between Circadian Time (CT) 1-3 for each cage of mice, where CT 0 was defined as 12 hours after the start of activity. Tests were performed in order of least to most stressful (i.e., open field, EPM, three-chamber social interaction, PPI, Barnes maze). Two days of rest were scheduled between each test.

### Open field test

We used the VersaMax Legacy Open Field setup (AccuScan Instruments, Inc., Columbus, OH, USA), and collected data using the Versamax Software (version 4.0, 2004; AccuScan Instruments, Inc.). Measures such as thigmotaxis (time spent in the outer edges divided by center) and total distance traveled and were analyzed(43).

### Elevated plus maze (EPM) test

The elevated apparatus consisted of a plus-shaped maze where 2 opposing arms (closed arms) were enclosed by high walls, while the other 2 opposing arms (open arms) had no walls. Mice were placed in the intersection of the four arms, facing an open arm. Mice freely explored the apparatus for 5 minutes. Time spent and entries in each arm was scored. See EPM formula below.

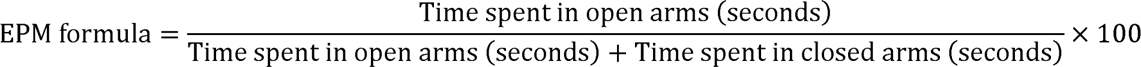

### Three-chamber social interaction test

Social preference and social memory were assessed using the three-chamber social interaction test(44). In the social preference phase, a mouse that our experimental mouse had never interacted with before, called stranger 1, was placed under one of the wire containers, and an object under the other wire container. See social preference formula below.

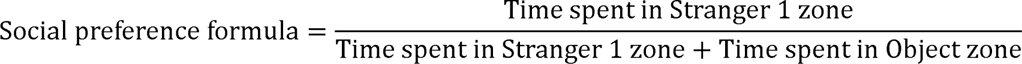

In the social memory phase, the object was replaced by a novel mouse, called stranger 2.

See social memory formula below.

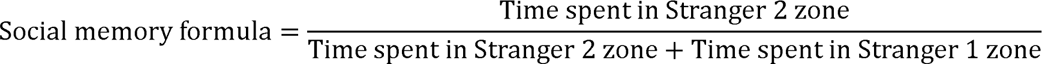

### Barnes maze

The Barnes maze was a circular platform (120 cm diameter) elevated 150 cm off the ground, with 20 holes around the perimeter. A camera and bright light were placed above the platform. A white noise sound was played (Play Store application called “white noise” on android, sound: blue noise, system volume: 10, 50 decibels) to motivate mice to seek shelter in an escape box, placed under one of the 20 holes around the apparatus. Mice were trained to locate the escape box using a configuration of visual cues located around the testing area (an image of a black square, circle and triangle located equidistant around the apparatus). There were two training days (the first had three trials and the second had two) where mice were given three minutes to locate the escape box. If they failed, mice were gently guided to the escape box, and a lid was placed on top for 1 minute. Latency to the escape box was measured for each training trial. On the test day, the escape box was removed, and mice had a total of 90 seconds to move freely on the apparatus. The duration in each quadrant, entries and duration in target quadrant, latency to target quadrant and number of errors were assessed.

### Prepulse inhibition of acoustic startle (PPI)

PPI was measured using SR-LAB startle chambers (San Diego Instruments, San Diego, CA, USA), and a software package by SR-LAB. Mice were placed into a cylindrical Plexiglass enclosure. Each session consisted of 50 trials, some of which only had a startle “tone” (broad-based noise burst), and some trials had a prepulse played before the startle tone. A piezoelectric accelerometer fixed to the Plexiglass base was used to detect and transduce motion resulting from the animal’s startle response. Each trial began with a 5-minute acclimation period. In the first 6 and final 6 trials, a ‘startle tone’ (120 dB for 50 ms, broad-band noise burst) was played in the absence of a prepulse (called ‘startle-only’ trials). In the next 38 trials, 8 trials were ‘startle-only’ trials, 5 trials had no stimulus, and 25 trials had the startle tone preceded by a prepulse stimulus (30 ms, broad-band burst) that was either 6, 9, 12 or 15 dB above the background noise and presented 100 ms before the startle pulse.

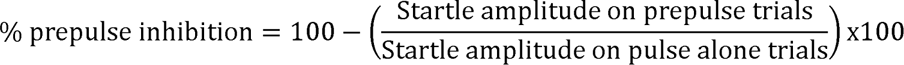

### Immunohistochemistry for microglial visualization

#### Microglia characterization

Microglia were characterized as described(35). Forty mice [5 mice per group: treatment (MIA, saline), lighting condition (LD, LL), sex (males, females)] were transcardially perfused using 4% paraformaldehyde (# 31985-062, Invitrogen). Sagittal brain sections (30 μm thick) were collected, containing the medial prefrontal cortex (mPFC) (bregma +2.46 mm until +1.94 mm) and dorsal hippocampus (−1.46 mm until −2.06 mm)(45). Sections were processed for immunofluorescence staining for a microglial marker Iba1 (ionized calcium binding adaptor molecule 1), using Rabbit anti-Iba1 (1:1000, # 019-19741, Cedarlane) as primary antibody and Alexa fluor 594 Donkey anti-rabbit IgG (1:1000, # 711-585-152, Cedarlane) as secondary antibody. Sections were counterstained with DAPI (1:20000, # D9542, Sigma).

#### Microglia visualization and analysis

Iba-1 immunostained microglia from the dorsal hippocampus (dentate gyrus and CA1) and mPFC were imaged using Z-stacks at a 20X magnification. Sections were visualized using an Olympus BX63 automated fluorescence microscope. Images were analyzed using Fiji ImageJ software, while being blinded to the experimental conditions. Measures of morphology included soma area and perimeter, arborization area and perimeter and soma circularity. Morphological index was calculated by dividing the soma area by the arborization area for each cell. The ‘nearest neighbor distance’ Fiji plugin was applied to determine the distance between each cell and its closest neighbor. The density of microglia for each image was calculated by dividing the number of cells per image by the area of the region of interest. The spacing index was calculated by multiplying the average density for each animal obtained above by the square of the average nearest neighbor distance.

### RNA sequencing

#### Tissue collection and pre-processing of data

The transcriptome was assessed from the dorsal hippocampus (1 mm thick punch containing bregma –1.46 mm until −2.46 mm) using next generation bulk RNA sequencing. Thirty-two mice [4 mice per group: treatment (MIA, saline), lighting condition (LD, LL) and sex (males, females)] were euthanized by decapitation, and brains were rapidly extracted and sliced (1 mm thick). The dorsal hippocampus was microdissected using a 0.5 mm diameter tissue punch and flash frozen on dry ice. RNA was extracted using Trizol/chloroform, and RNA quantity and integrity was analyzed using a Nanodrop (Thermo Fisher). Sequencing was performed at the IRIC (Institute for Research in Immunology and Cancer) genomics sequencing platform at the University of Montreal (Nextseq 1×75bp, around 20M reads/samples). Pre-alignment quality control was performed at the sequencing platform using FastQC. Alignment was performed using the STAR version 2.7.1a aligner to the GRCm38 / mm10 Mus musculus genome assembly (<u>Mus musculus genome assembly GRCm38 - NCBI - NLM</u> (nih.gov). Genes that did not meet the requirement of a minimum of one count per million (cpm) in at least 4 samples were excluded. Only genes annotated as protein coding according to the ensembl’s biomart Mus musculus package were used for analyses (https://bioconductor.org/packages/release/data/annotation/html/Mus.musculus.html). Outliers were assessed in normalized data using unsupervised hierarchical clustering of mice by principal component analysis. RNA-sequencing data were deposited in the Gene Expression Omnibus database and will become available upon publication of the peer-reviewed manuscript.

#### Differential expression (DE) analysis

DE analysis was performed using Deseq2 package in R(46), which modelled the raw counts (using normalization factors to account for differences in library depth) and estimated the gene-wise dispersions and shrunk these estimates to generate more accurate estimates of dispersion to model the counts. Then, Deseq2 fit the negative binomial generalized linear model.

#### Rank rank hypergeometric overlap 2 algorithm (RRHO2)

RRHO2 is a threshold-free algorithm used to reveal overlapping trends in gene expression datasets(47). Differential expression lists were ranked by their −log_10_(p-value) and effect size direction, then lists were compared to identify significant overlap in genes across a continuous significance gradient. Concordant and discordant gene overlap (assessed for sex differences, and the effects of LL and MIA) described genes that were respectively up and downregulated in both gene expression signatures, as well as overlap of expression profiles in opposite directions(47). RRHO maps were used to visualize the intensity of significant overlap between two ranked lists. They computed the normal approximation of difference in log odds ratios and standard error of overlap. The z-scores were then converted to a p-value and corrected for multiple comparisons across pixels.

#### Weighted Gene Co-expression network analysis (WGCNA)

The WGCNA R package constructs a signed matrix representing pairwise correlations among all gene pairs across the selected samples(48). The soft-threshold power was determined based on gene-to-gene connections and adhered to the scale-free topology criterion. Following this, the correlation matrix was then transformed into an adjacency matrix, with genes represented as nodes and the connection strength between genes as edges. Subsequently, this matrix was transformed into a topological overlap matrix (TOM), to divide the genes into distinct modules (clusters) of co-expressed genes. The dynamic tree cut algorithm was used to divide the hierarchical clusters into gene modules, ensuring a minimum module size of 30 genes. Module-trait associations were determined by calculating the correlation between the module eigengene (ME) and a condition (MIA exposure and LL exposure) for each module. A correlation coefficient (r) and p-value were reported for each identified module. Modules with statistically significant correlations (p<0.05) with either MIA exposure or LL exposure were further analyzed. We used automatic network construction and module detection functions (blockwise) with the following parameters: signed network; Pearson correlation; maximum Block Size of 7000 genes; merge Cut Height of 0.25.

#### Identification of hub-bottleneck genes and pathway enrichment analysis

The topological properties of the gene network were calculated using the CytoscapeSoftware® data. As previously described, ‘degree’ was calculated by summing the number of adjacent nodes, which describes the local topology, and ‘betweeness’ considers the number of shortest paths linking two nodes and passing through a node n(49). Nodes with ‘degree’ values higher than +1 SD above the mean were considered hubs, and nodes with ‘betweenness’ values higher than +1 SD above the mean were considered bottlenecks. Nodes considered both hubs and bottlenecks (hub-bottlenecks) were identified as central nodes within networks(50, 51). An enrichment analysis was then performed using the entire mouse genome as the background gene set. We used functional enrichment analysis (MetaCore®, Clarivate Analytics, https://portal.genego.com) to investigate significant gene ontology processes associated with central nodes within networks.

#### Overlap between hub-bottleneck genes and genes related to sleep/circadian rhythms and neurodevelopmental disorders

Hub-bottleneck genes from the mouse dataset were first converted to their human homologs (using the Bioconductor package in R BiomaRt, with dataset = “hsapiens_gene_ensembl and dataset = “mmusculus_gene_ensembl, https://dec2021.archive.ensembl.org). We identified genes that were common to both the list of hub-bottleneck genes and those associated with sleep/circadian rhythms and neurodevelopmental disorders as defined by genome-wide association studies (GWAS). The GWAS catalog (https://www.ebi.ac.uk/gwas/home), which compiles available GWA studies and offers clearly searchable SNP-trait associations, was searched for traits of interest. The following datasets were used: 8 datasets associated with ‘sleep’ (trait labels: sleep depth EFO_0005273, sleep efficiency EFO_0803364, sleep latency EFO_0005280, sleep quality EFO_0005272, sleep measurement EFO_0004870, sleep time EFO_0005274, sleep duration EFO_0005271, sleep disorder EFO_0008568), 3 datasets associated with ‘circadian rhythm’ (trait label: circadian rhythms, including disrupted circadian rhythms, i.e., low relative amplitude of rest-activity cycles EFO_0004354, publication exploring chronotype PubMedID 30696823, publication exploring actigraphy PubMedID 33075057), 1 dataset associated with ‘microglia activation’ (trait label: microglial activation measurements EFO_0010940), 4 datasets associated with ‘autism spectrum disorders’ (trait labels: autism spectrum disorder EFO_0003756, autism EFO_0003758, autism spectrum disorder symptom EFO_0005426, and asperger syndrome EFO_0003757), and 3 datasets associated with ‘schizophrenia’ (trait labels: schizophrenia MONDO_0005090, treatment refractory schizophrenia EFO_0004609, and schizophrenia symptom severity measurement EFO_0007927).

#### Cell-type specific gene set enrichment within WGCNA modules and among differentially expressed genes

Gene set enrichment analysis (GSEA) was used to determine the genes in the WGCNA modules of interest that are overexpressed within differentially expressed genes (DEGs) lists associated with LL or MIA. We performed GSEA using the fgsea package (v1.28.0) on R (v4.3.2) of the brown, purple and cyan modules on the LL DEG list, and the brown, purple and green modules on the MIA DEG list. For each module, we selected the leading edge (LE) genes and their p-values in the LL and MIA DEG lists for Differential cell type change analysis using Logistic Regression (LRcell) (Ma et al., 2022). The LRcell package (v1.10.0) (52)was used to determine the cell types driving the differential expression of the selected genes using the cell type marker genes from the mouse hippocampus integrated in the LRcell package from Saunders and colleagues(53). The results are represented as Manhattan plots with the cell types on the x-axis and the negative logarithmic transformation of the FDR, with at a significant cut-off of 0.05. Here, we only report LRcell tests with significant results. The LRcell analysis was also applied to the complete LL and MIA DEGs lists. Only the results for modules and DEG lists that yielded a significant enrichment of at least one cell type are reported.

#### Expression patterns of WGCNA module genes in a single cell RNA sequencing dataset

We used the CZ CELLxGENE gene expression tool (54) to create dot plots of genes from WGCNA modules of interest across cell types from the pre-processed and normalized single-cell data from a published study(55). All hub-bottleneck genes of the brown, purple, cyan and green modules were used. For the cyan and green modules, the analysis was also done with the entire list of genes. For each gene, the color of the dots represents the average gene expression level (normalized by logarithmic transformation of pseudo counts per 10,000 reads), scaled from 0-1 across the cell types select for the dot plot. The size of the dots represents the percentage of cells within each cell type that express that gene.

#### Statistics

Data were analyzed and graphed using Prism version 9 (GraphPad). Data describing number of pups per litter, and maternal cytokine and chemokine expression levels were analyzed using independent sample t-tests. Data that did not pass the Shapiro-Wilk normality test were analyzed by Mann-Whitney U test, and data with unequal variances (assessed by an F-test) were analyzed by an independent samples t-test with Welsh’s correction (Supplementary Fig. 1A, D, G; Supplementary Fig. 2). For offspring body weight data, we conducted 2-way mixed effect ANOVAs, with treatment (poly IC, saline) as a between-subjects variable and age (post-natal week 3, 5, 8) as a within-subjects variable. We explored treatment x age interactions and main effects for each sex separately (Supplementary Fig. 1B, C, E, F, H, I). For the behavior and microglia data, 2-way mixed effect ANOVAs were conducted using Tukey’s post hoc comparisons, with treatment (poly IC, saline) and lighting condition (LD, LL) as a between-subjects variables. Main effects were explored if no significant group x lighting interaction was observed (Fig. 2-6, Supplementary Fig. 3-6). Three-way ANOVAs were also conducted (using SPSS statistical software version 29) for prepulse inhibition (3^rd^ factor: prepulse) and locomotion (3^rd^ factor: time) (Supplementary Fig. 3-4). WGCNA (Fig. 7, 8) and RRHO (Supplementary Fig. 8) analyses were assessed as previously described(47, 48). Sex differences (Supplementary Tables 1, 2) were assessed using three-way ANOVAs (treatment x lighting x sex) and decomposed by assessing the 2-way interaction (treatment x sex) and observing the main effect of sex. Differences were considered significant if *p* < 0.05.

**Fig. 2.**
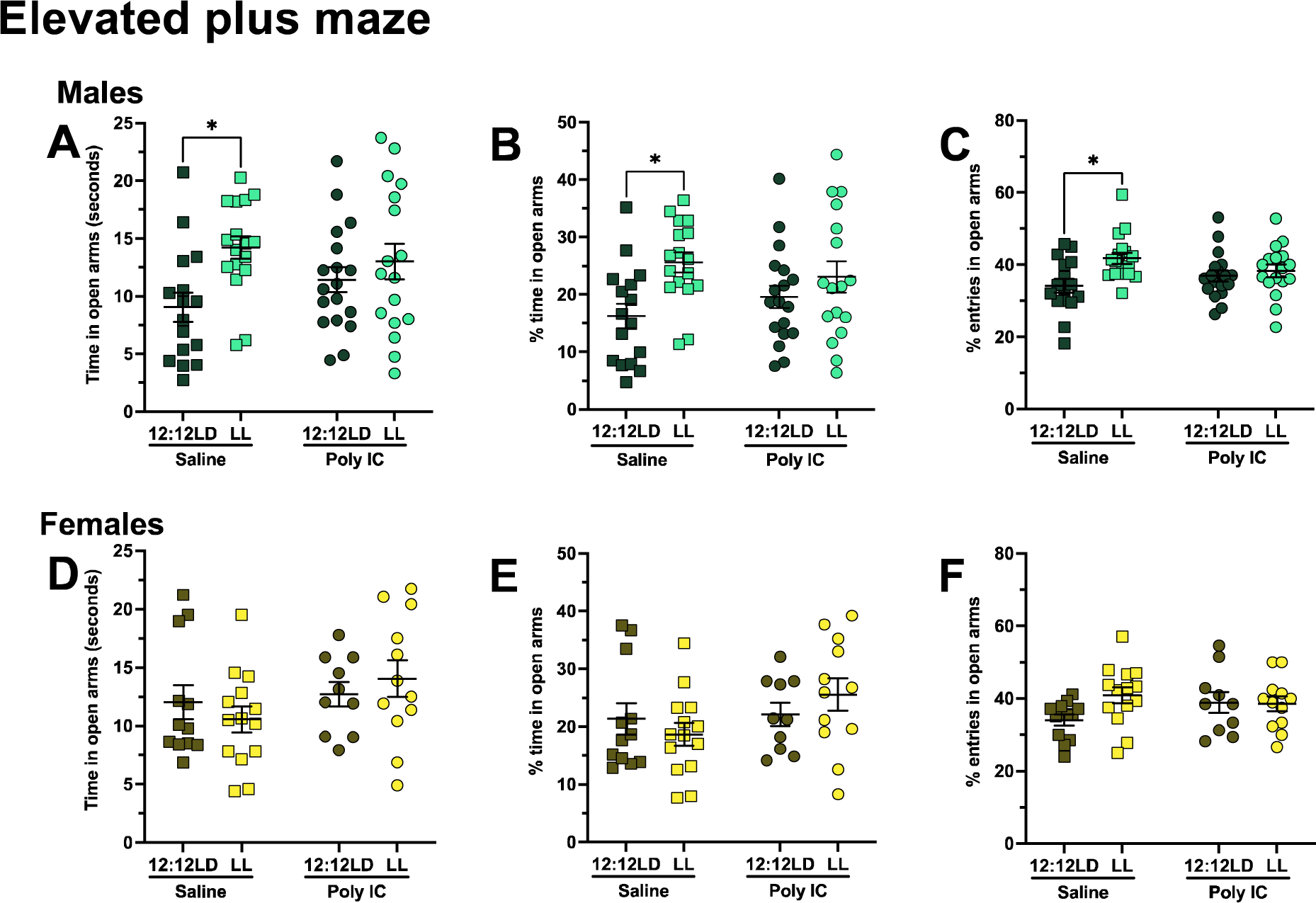
LL leads to decreased anxiety-like behavior in saline-but not poly IC-exposed male mice and not in females. Time in open arms (**A, D**), percent time in open arms (**B, E**) and percent entries in open arms (**C, F**) were assessed in males (**A-C**) and females (**D-F**). Data points represent individual mice, and are presented as mean ± SEM. Two-way ANOVAs (factors treatment x lighting with Tukey’s post-hoc comparisons) were conducted. \**p* < 0.05 (post hoc).

**Fig. 3.**
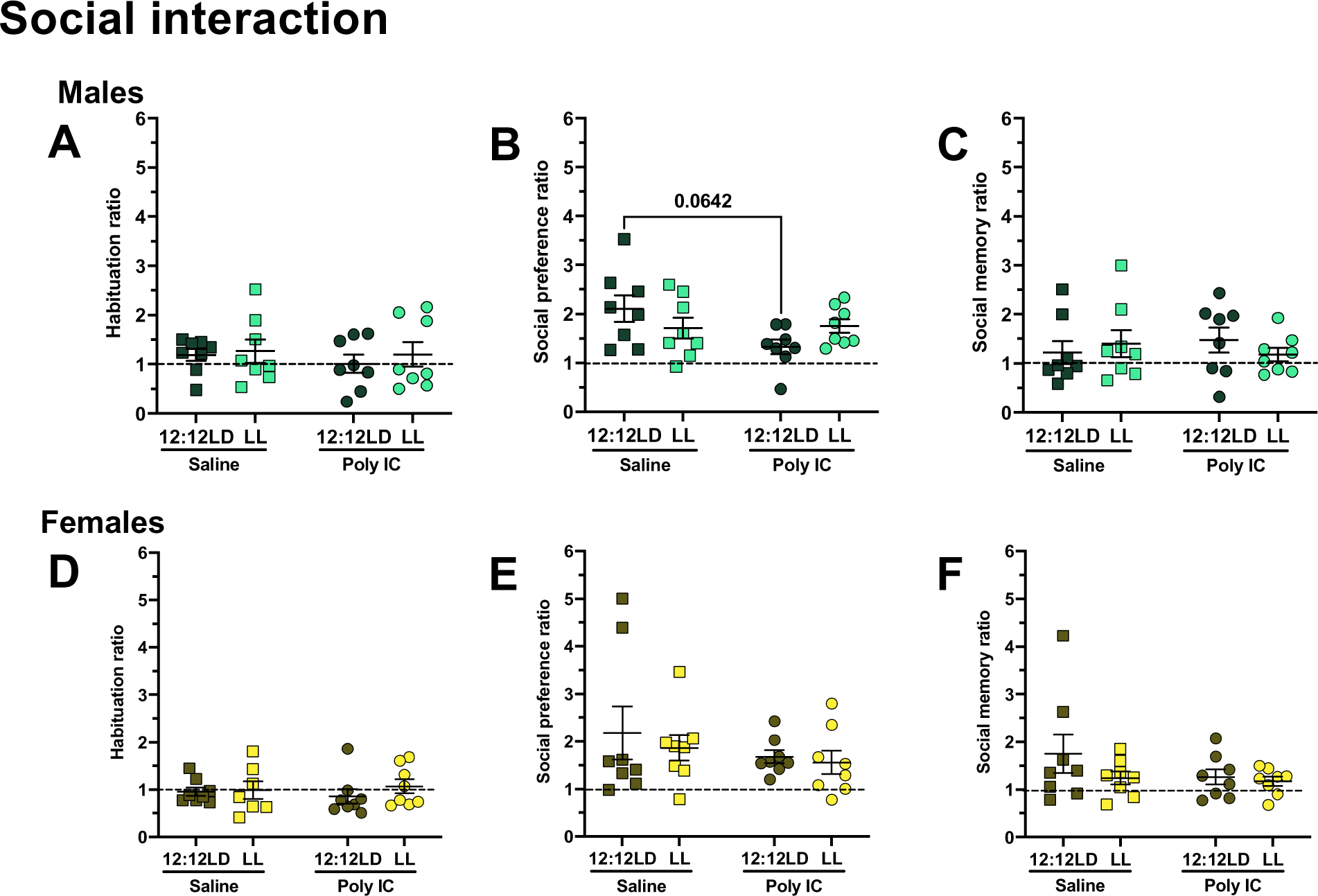
Reduced sociability in poly IC-exposed males after 12:12LD, but not LL. Preference ratios were assessed for the habituation **(A, D)**, sociability (**B, E**) and social memory phases (**C, F**) in males (**A-C**) and females (**D-F**). Data points represent individual mice, and are presented as mean ± SEM. Two-way ANOVAs (factors treatment x lighting with Tukey’s post-hoc comparisons) were conducted.

**Fig. 4.**
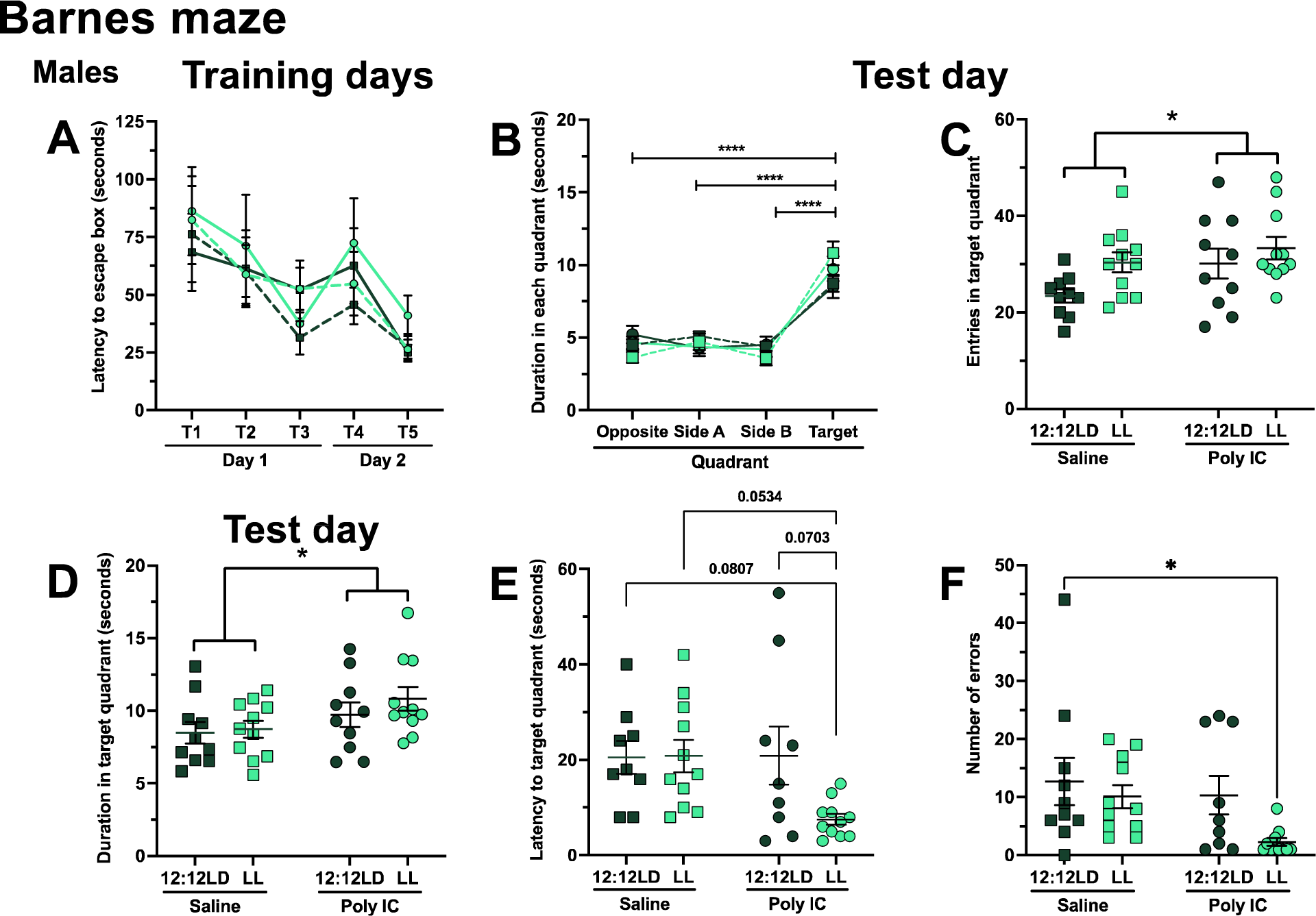
Poly IC exposure leads to improved spatial memory in males. Latency to escape box (**A**) was assessed for the training day. On the test day, duration in each quadrant (**B**), entries in target quadrant (**C**), duration in target quadrant (**D**), latency to target quadrant (**E**) and number of errors (**F**) was assessed. Data points represent individual mice, and are presented as mean ± SEM. A two-way ANOVA (factors group x trials) was conducted for the training days, another two-way ANOVA (factors group x quadrant) was conducted for duration in each quadrant, and additional two-way ANOVAs (factors treatment x lighting with Tukey’s post-hoc comparisons) were conducted for the remaining graphs. \**p* < 0.05, ****p<0.0001.

**Fig. 5.**
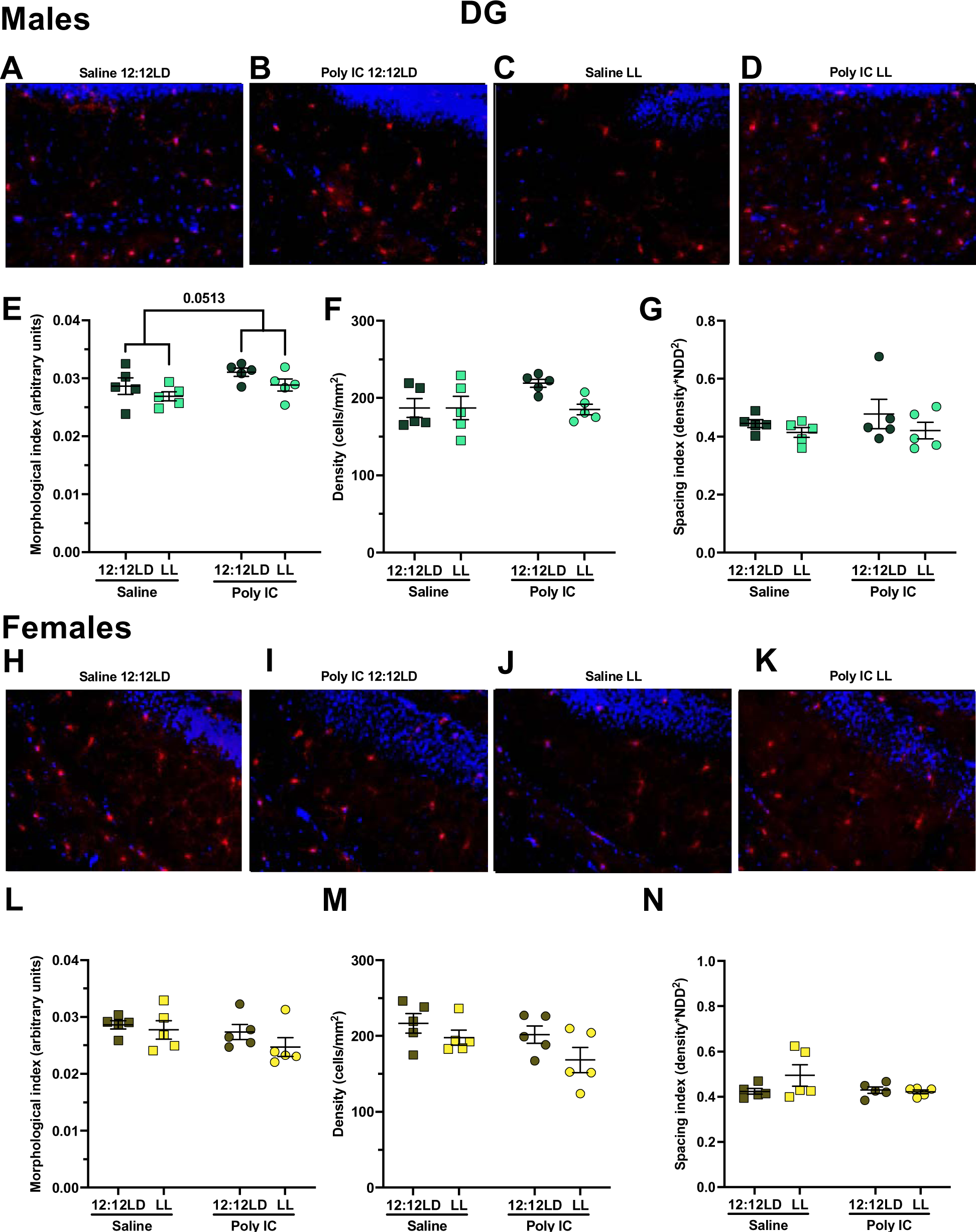
Poly IC exposure leads to increased morphological index but not density in the dentate gyrus. Representative images of microglia from the dentate gyrus for each group under each lighting condition are shown for males **(A-D)** and females **(H-K)**. Morphological index (**E, L**), density (**F, M**) and spacing index (**G, N**) were assessed for males (**E-G**) and females (**L-N**). Data points represent individual mice and are presented as mean ± SEM. Two-way ANOVAs (factors treatment x lighting with Tukey’s post-hoc comparisons) were conducted.

**Fig. 6.**
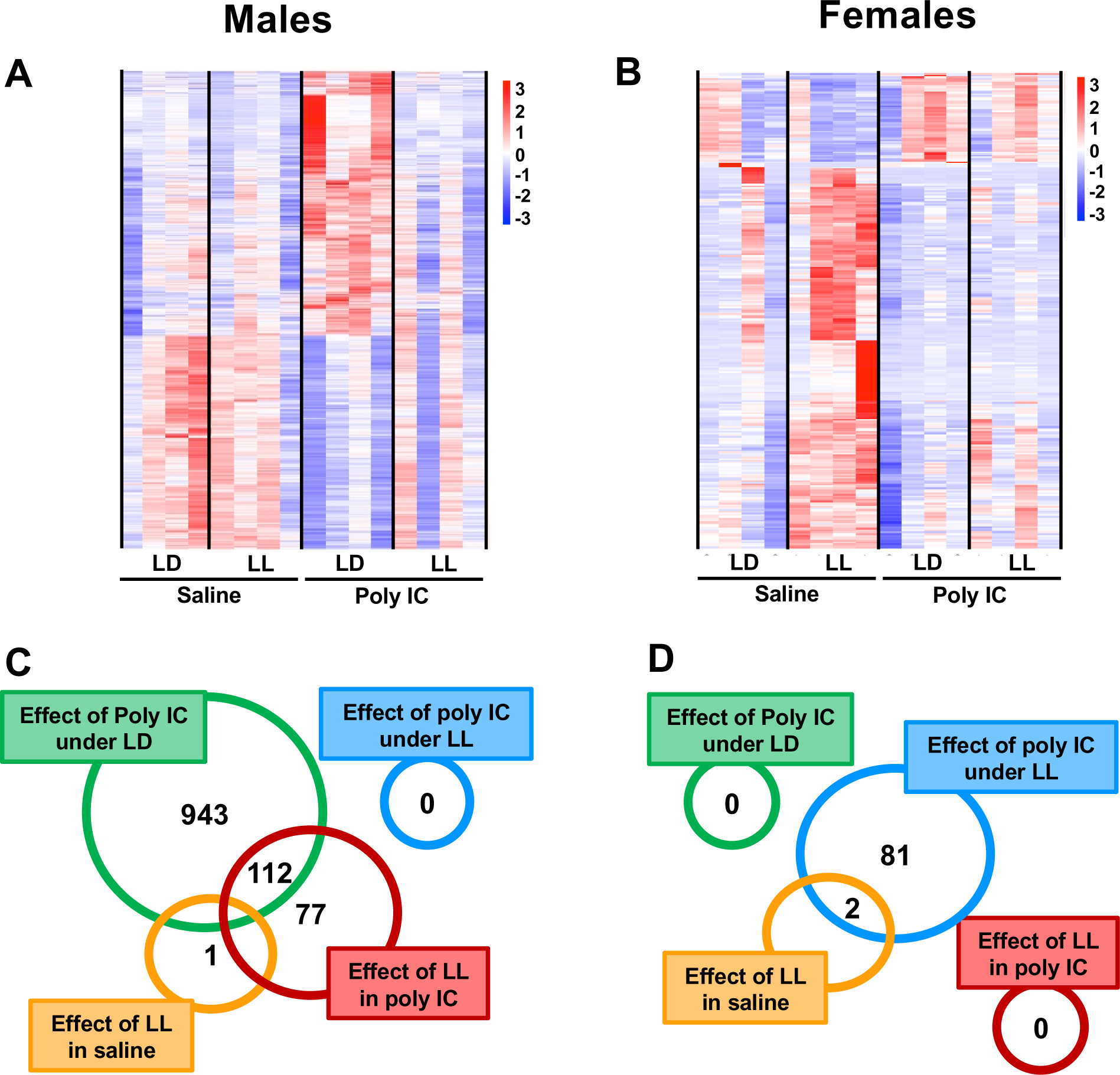
Sex-specific effects of poly IC and LL on the hippocampus transcriptome. Venn diagrams and heatmaps of all differentially expressed (DE) transcripts are shown for males (**A, C**) and females (**B, D**). DE analysis was performed using Deseq2 package and images were created using Rstudio®.

**Fig. 7.**
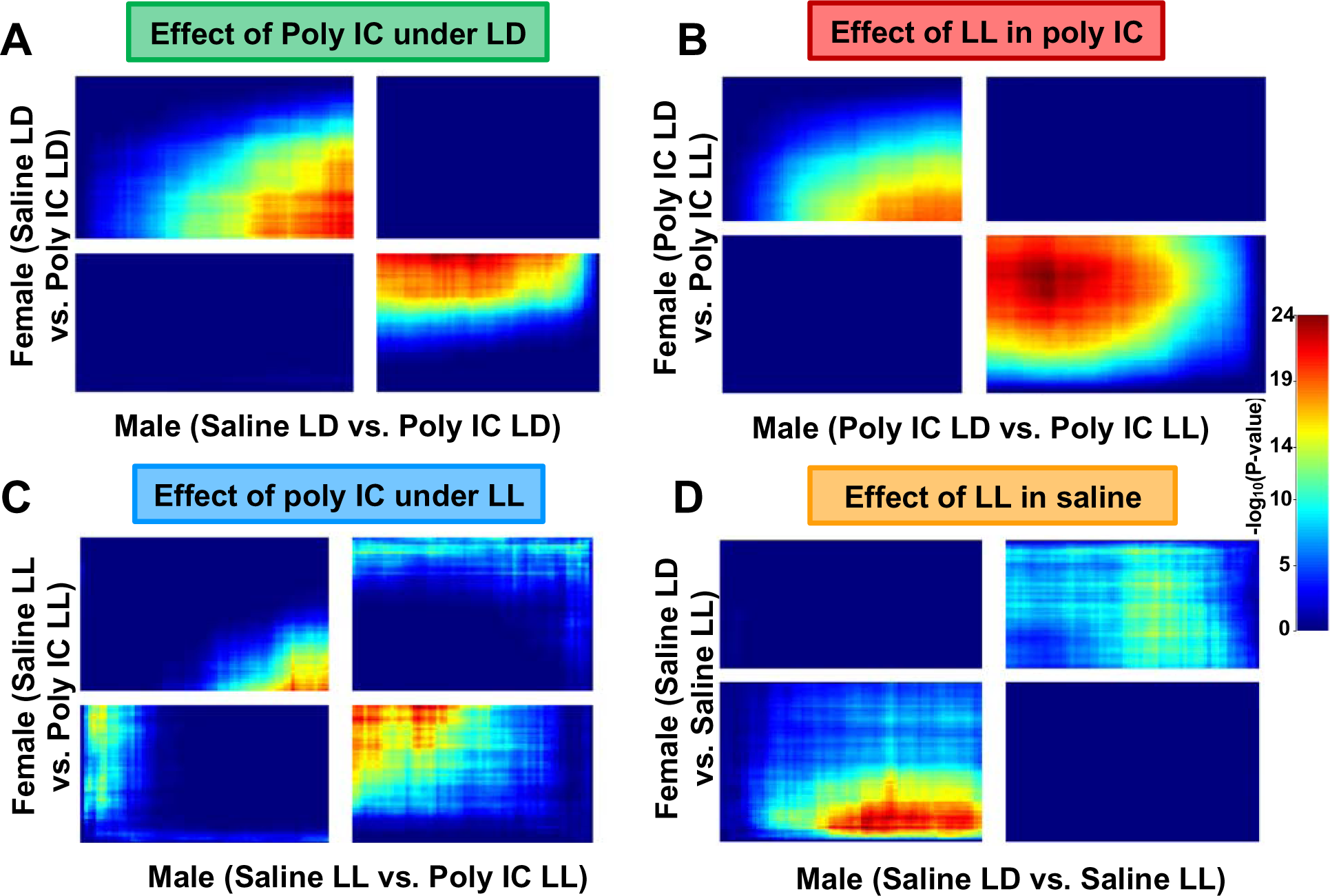
Rank rank hypergeometric overlap 2 (RRHO2) analysis highlights in discordant patterns between sexes. RRHO2 plot indicating discordance patterns between sexes, highlighting sex differences.

## Results

### Litter details

The litters of poly IC- and saline-exposed mice did not differ in the number of pups in any of the cohorts (cohorts used for: behavior, *p* = 0.9094; microglial characterization, *p* = 0.5009; RNA sequencing, *p* = 0.3527) (Supplementary Fig. 1A, D, G). Only litters with 4 to 10 pups inclusively were used for testing; 1 saline and 2 poly IC from the first cohort (behavior) were excluded as these litters only had 3 pups. Mice from 4-6 different litters were used in each experimental condition, and we visually confirmed that significant effects reported were not driven by litter effects. Poly IC-exposed offspring weighed significantly less than saline-exposed offspring in the cohort used for behavior (Supplementary Fig. 1B, C) (effect of treatment male: p = 0.0006, female: p < 0.0001), but not the cohorts used for microglia analysis or transcriptomics (Supplementary Fig. 1E, F, H, I).

### Poly IC induced inflammatory response in pregnant dams

Poly IC-injected dams had a significant elevation in blood plasma expression of the following pro- and anti-inflammatory cytokines: IL-6 (*p* = 0.0427), IL-10 (*p* = 0.0014), TNFα (*p* = 0.0029) and MCP-1 (*p* = 0.0384) compared to controls, with no significant differences in IL-12(p70) (*p* = 0.4608), IL-2 (*p* = 0.8754), IL-1β (*p* = 0.6798), IL-4 (*p* = 0.4990) and GM-CSF (*p* = 0.2844) (Supplementary Fig. 2). These data confirm that the poly IC lot used in the experiments was able to generate an immune response in the dams.

### Behavior

#### Similar spontaneous locomotion in saline and poly IC-exposed mice

Mice from all experimental groups had similar scores on total distance traveled in the open field test (Supplementary Fig. 3A, C).

#### Decreased anxiety-like behavior after LL in saline-but not poly IC-exposed male mice

While no significant interactions were observed in males, significant main effects of lighting were noted in time spent in open arms (*p* = 0.0078), percent time in open arms (*p* = 0.0045) and percent entries in open arms (*p* = 0.0091). Post hoc tests revealed that saline/LL males had decreased anxiety-like behavior compared to saline/LD, as reflected by more time spent in open arms (*p* = 0.0204), percent time in open arms (*p* = 0.0237) and percent entries in open arms (*p* = 0.0121). Interestingly, LL did not have this effect in poly IC-exposed males (Fig. 2A-C). Females showed no significant differences in tested parameters (Fig. 2D-F). Of note though, differences were observed in thigmotaxis ratio in the open field test (Supplementary Fig. 3B, D). In conclusion, LL exposure led to decreased anxiety-like behavior in saline-but not poly IC-exposed males.

#### Reduced sociability in poly IC-exposed males after 12:12LD, but not LL

During the habituation phase of the three-chamber social interaction test, mice spent a similar amount of time in each chamber (Fig. 3A, D). Poly IC-exposed males exhibited a trend for decreased sociability (treatment x lighting interaction: *p* = 0.0508), specifically under 12:12LD (post hoc: *p* = 0.0642), but not LL (Fig. 3B). No differences were observed between groups in females (Fig. 3E) and no significant differences were observed in social memory in either males or females (Fig. 3C, F). Altogether, poly IC exposure led to a trend for reduced sociability in males under 12:12LD but not LL exposure and these effects were not seen in females.

#### Poly IC exposure led to improved spatial memory in males with apparent modulation by LL

During the training phase of the Barnes maze test, all groups spent a similar amount of time locating the escape box (Fig. 4A). On the test day, all groups spent significantly more time in the target quadrant compared to other quadrants (quadrant x treatment interaction: p = 0.0427; post hocs between target quadrant and each of the other quadrants, *p* < 0.0001) (Fig. 4B). Poly IC exposure led to increased entries (*p* = 0.0438) and duration (*p* = 0.0322) in the target quadrant, without significant treatment x lighting interactions (Fig. 4C, D). A trend for a group x lighting interaction was observed in the latency to reach the target quadrant (*p* = 0.0752), with post hoc tests revealing a trend for a difference between poly IC/LL and all other groups: saline/12:12LD (*p* = 0.0807), poly IC/12:12LD (*p* = 0.0703) and saline/LL (*p* = 0.0534) (Fig. 4E). Although no significant treatment x lighting interaction was found, poly IC-exposed males seemed to make less errors to the target (effect of treatment, *p* = 0.0679) and the poly IC/LL group may be driving these effects (post hocs: saline/12:12LD vs. poly IC/LL*, p* = 0.0422) (Fig. 4F). Overall, these data show that poly IC-exposed males performed better on the Barnes maze compared to saline-exposed males, especially the poly IC/LL group.

#### No sensory motor gating deficits following poly IC exposure

In the PPI test, no significant differences were observed in males and females in baseline startle response, average percent prepulse inhibition or percent prepulse inhibition over time (Supplementary Fig. 4).

#### Sex differences

Sex differences were assessed in the behavioral data, and no treatment x sex x lighting or treatment x sex interactions were found in any of the tested behavioral parameters. We report a main effect of sex in the thigmotaxis ratio in the open field test, where females had significant higher scores than males, indicative of more anxiety-like behavior (p<0.001). Additionally, we found a significant main effect of sex for baseline startle response in the PPI test where females had significantly lower scores than males (*p* = 0.0485) (Supplementary Table 1). Sex differences were also assessed in the microglia data and while no treatment x sex x lighting interactions were found, we did report some treatment x sex interactions. Namely, in the dentate gyrus, we found a significant treatment x sex interaction for morphological index (*p* = 0.0183) and density (*p* = 0.0320). Lastly, we found several main effects of sex, including on morphological index in the dentate gyrus (*p* = 0.0496) and prefrontal cortex (*p* = 0.0160) (Supplementary Table 2).

#### Poly IC exposure led to an increased activated profile of microglia in the dentate gyrus

Microglia were characterized in male and female mice exposed to in utero poly IC or saline. In the dentate gyrus, poly IC exposure led to significantly increased morphological index compared to saline-exposed males but not females (main effect of treatment, *p* = 0.0513), with no significant treatment x lighting interaction (Fig. 5D, J), and no differences in microglial density or spacing index (Fig. 5E-F, F-L). No differences were observed in the prefrontal cortex or CA1 for either sex (Supplementary Fig. 5-6). Overall, differences were only observed in the dentate gyrus, with microglia of poly IC-exposed males exhibiting a more inflammatory morphological phenotype.

### RNA sequencing

#### Sex-specific effects of poly IC and LL on the hippocampus transcriptome

RNA sequencing was performed on hippocampal tissue of saline- and poly IC-exposed males and females. Heatmaps represent all differentially expressed (DE) transcripts in the hippocampus for males and females, whereby each row represents a gene, and each column represents a mouse (Fig. 6A, B, Supplementary Files 1, 2). A high number of DE transcripts were present in poly IC-exposed males (1055) compared to saline-exposed males, with an almost even distribution of downregulated (590) and upregulated (465) transcripts (Fig. 6A, C). In females, few genes were DE between groups, except for 83 genes being DE between saline- and poly IC-exposed mice under LL (70 downregulated and 13 upregulated transcripts) (Fig. 6B, D). Surprisingly, in both sexes, there were very few DE transcripts between saline-exposed mice after LL exposure and saline-exposed mice after LD (male = 1 downregulated transcript, females = 1 upregulated and 1 downregulated transcripts). However, in poly IC-exposed males (but not females), LL exposure led to 189 DE transcripts (110 upregulated vs. 79 downregulated transcripts) in comparison to poly IC-exposed males after LD exposure (Fig. 6C). Overall, poly IC exposure seems to affect the male hippocampal transcriptome more than the female transcriptome, and poly IC-exposed males seem sensitive to LL exposure, while females do not.

LRcell analysis did not reveal any significant cell types associated with differential gene expression after MIA exposure. However, many cell types reached significance in the LRcell analysis using the LL DEGs list (Supplementary Fig. 7), with a neurogenesis neuron cell type enriched for *Sox4* and *Gabra5* (HC_13.5) being the most significant. All but one of the oligodendrocyte subtype clusters reached significance, indicating that these cells are implicated in the changes in gene expression after LL exposure.

#### Rank-rank hypergeometric overlap (RRHO) analysis results in discordant patterns between sexes

We explored the extent of transcriptional overlap between sexes using RRHO2, which orders transcripts by effect size direction and p value. The bottom left and top right quadrants in RRHO2 maps represent regions of concordant overlap between differential expression directions, with the top left and bottom right regions being discordant. We found discordant patterns of expression when comparing sex differences in: the effect of poly IC under LD (saline LD versus poly IC LD) (Fig. 7A), the effect of LL in poly IC-exposed mice (poly IC LD versus poly IC LL) (Fig. 7B), the effect poly IC under LL (saline LL versus poly IC LL) (Fig. 7C) and the effect of LL in saline-exposed mice (saline LD versus saline LL) (Fig. 7D). In addition, in both males and females, we found substantial overlap in transcripts when comparing the effect of LL in poly IC-exposed mice (poly IC LD versus poly IC LL) and saline-exposed mice (saline LD versus saline LL) (Supplementary Fig. 8A, B), and when comparing the effect of poly IC between mice exposed to LD (saline LD versus poly IC LD) and those exposed to LL (saline LL versus poly IC LL) (Supplementary Fig. 8C, D).

#### Modules of co-expressed genes associated with poly IC and LL exposure in males and females

Gene expression from male and female mouse hippocampus was analyzed using WGCNA to identify modules of co-expressed genes associated with either poly IC or LL exposure (Figure 8, Supplementary Files 3, 4). WGCNA identified a total of 44 modules (networks of co-expressed genes) in males and 26 in females. Using the module eigengene-trait analysis, we found that 4 of the 44 modules in males showed statistically significant associations with poly IC treatment. Specifically, 1 module (green, *p* = 0.04) was positively correlated and 3 modules were negative correlations (turquoise, *p* = 0.03; saddle brown, *p* = 0.02; purple, *p* = 0.02) with poly IC treatment. Two additional modules (brown and light green) showed a trend for a negative correlation (*p* = 0.06 and 0.05, respectively). Further, 8 of the 44 modules in males showed statistically significant associations with LL exposure. Specifically, 4 showed positive correlations (purple, *p* = 0.01; cyan, *p* = 0.02; skyblue3, *p* = 0.003; brown, *p* = 0.02; a fifth module, white, shows a trend for a positive correlation, *p* = 0.05), and 3 showed negative correlations (dark grey, *p* = 0.04; magenta, *p* = 0.00002; darkgreen, *p* = 0.01) with LL. Interestingly, some modules were associated with both poly IC treatment and LL exposure in males. For example, the purple showed a statistically significant positive correlations with LL exposure, and a statistically significant negative correlation with poly IC treatment (the brown module had a significant positive correlation with LL, and a trend for a negative correlation with poly IC) (Fig. 8A). Additionally, 4 of the 26 modules in females showed statistically significant associations with poly IC treatment, with 3 modules being negatively correlated (light green; *p* = 0.007; magenta, *p* = 0.02; cyan, *p* = 0.02) and 1 module being positively correlated (royal blue, *p* = 0.009). In females, no modules were significantly associated with LL exposure; however, 2 modules had a high correlation coefficient (turquoise, 0.49, *p* = 0.06 and blue, −0.46, *p* = 0.07) (Fig. 8B). Overall, more modules were significantly associated with poly IC or LL exposure in males than in females.

**Fig. 8.**
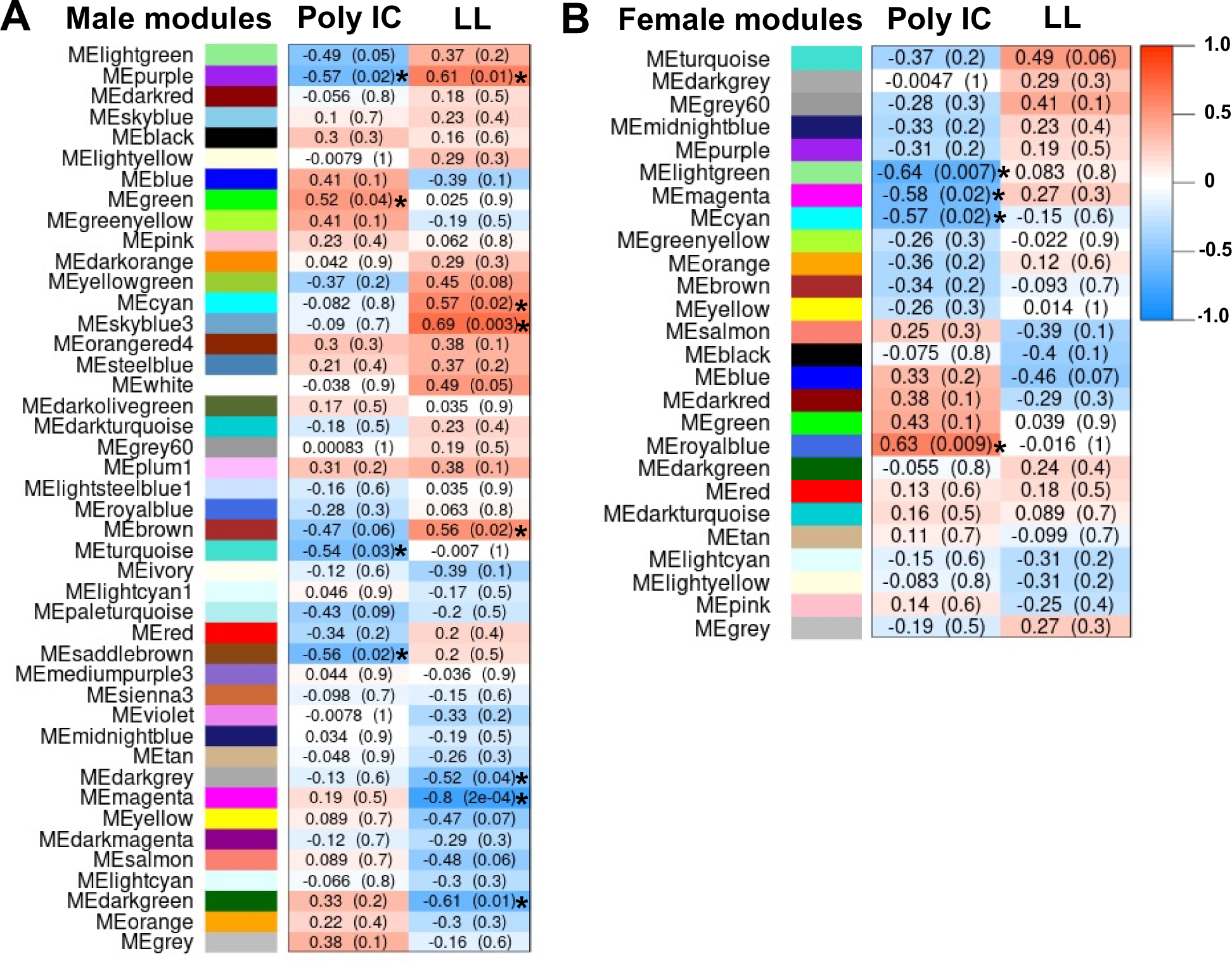
Significant modules associated with poly IC and LL exposure in males and females. Weight gene correlation network analysis (WGCNA) was used to generate gene co-expression modules from each condition separately (treatment, lighting). A correlation coefficient (r) and p-value are reported for each identified module. Modules significantly associated with poly IC or LL exposure are indicated with an asterisk.

#### Association of LL or poly IC hub-bottleneck genes with synaptic signaling pathways and with genes related to sleep/circadian rhythms and neurodevelopmental disorders in humans

We then identified the hub-bottleneck genes of the WGCNA modules, i.e. the genes acting as central nodes within the networks (Supplementary File 5). In males, the hub-bottleneck genes from the brown (144 genes) and purple (23 genes) modules (positively associated with LL and negatively associated with poly IC) were enriched for pathways involved in the modulation of synaptic transmission and vesicle mediated transport, as determined by Metacore (Fig. 9A, B). Genes that were central to the brown and purples modules (hub-bottleneck genes) include *Prelid3a*, which is involved in phospholipid transport, *Tmed9*, which is part of a family of genes encoding transport proteins in the endoplasmic reticulum and the Golgi, and *Snca*, which is found primarily in presynaptic terminals, playing a role in the transport of synaptic vesicles and which has been implicated in Parkinson’s disease (Fig. 9A, B). Additionally, many genes in these modules, including hub-bottleneck genes, were homologous to human genes identified in GWAS for sleep/circadian rhythms (*Prr7* and *Map2k1)* or schizophrenia (*Arhgap15, Snca, Erc2, Lrrtm4* and *Rbfox1*).

**Fig. 9.**
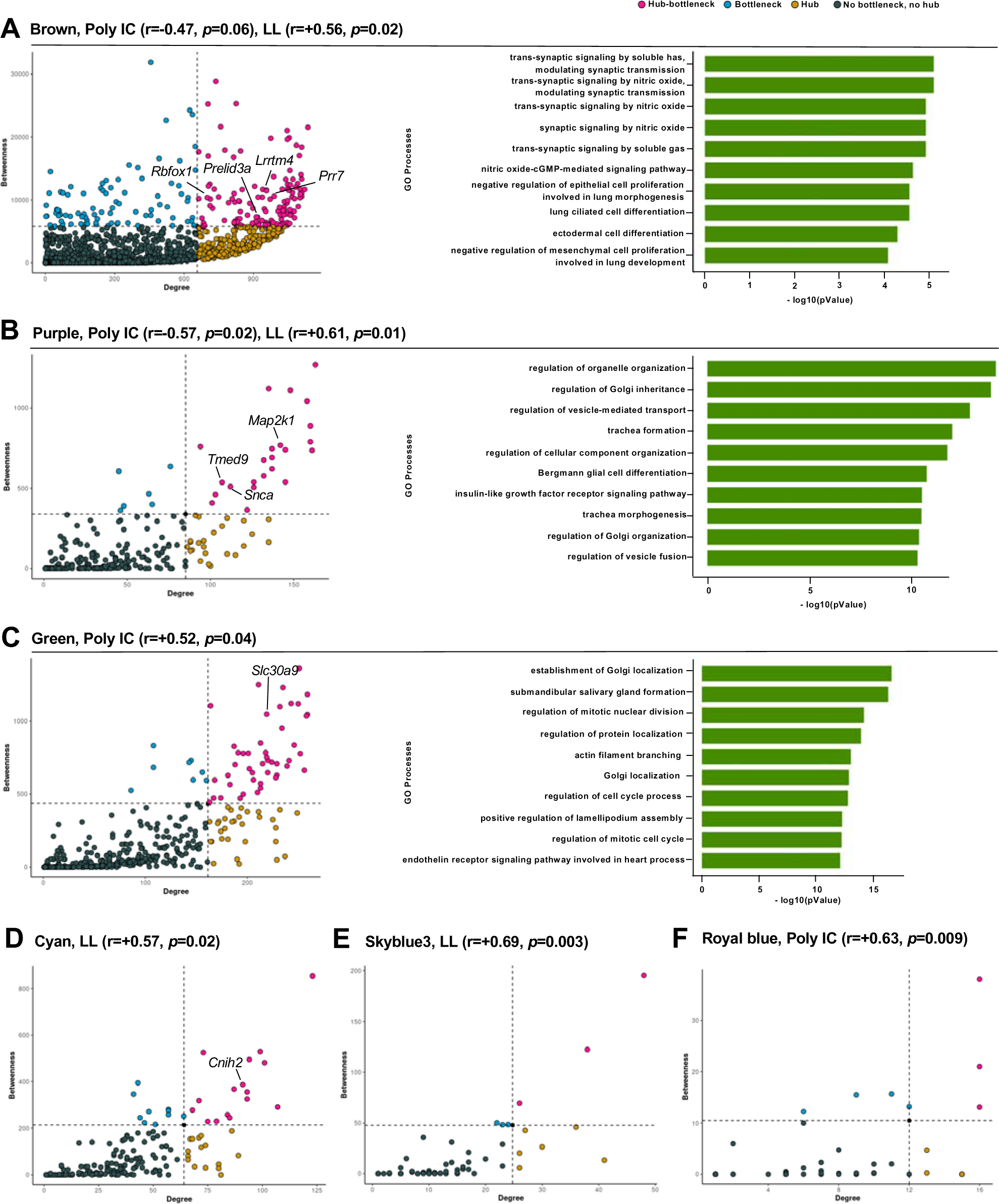
Hub-bottleneck genes show enrichment in pathways associated with augmented synaptic signalling in the brain. CytoscapeSoftware® was used to calculate topological properties of the gene networks. ‘Degree’ was calculated by summing the number of adjacent nodes, and ‘betweeness’ considers the number of shortest paths linking two nodes and passing through a node. Nodes with ‘degree’ values higher than +1 SD above the mean were considered hubs, and nodes with ‘betweenness’ values higher than +1 SD above the mean were considered bottlenecks. Nodes considered both hubs and bottlenecks (hub-bottlenecks) were identified as central nodes within networks. Functional enrichment analysis (MetaCore®, Clarivate Analytics) was used to investigate significant gene ontology processes associated with central nodes within networks. Selected hub-bottleneck genes indicated in the graphs are discussed in the text.

Also in males, hub-bottleneck genes from the green (46 genes) module (positively associated with poly IC) were enriched (Metacore) for pathways involved in cellular metabolic processes (Fig. 9C). Many genes in these modules, including hub-bottleneck genes, were homologous to human genes identified in GWAS for ASD and SCZ (*Slc30A9, Znf195* and *Gabbr1*), but also genes related to sleep (*Shb, Arhgap11a*).

In the cyan (16 genes) module (positively associated with LL), *Cnih2,* a hub-bottleneck gene highly connected within the module, encodes a subunit of ionotropic glutamate receptor of the AMPA subtype (Fig. 9D). In this module, 4 genes from the family of chemokine (C-C motif) ligand (*Ccl23*, *Ccl15*, *Ccll15, Ccl14*) were homologous to human genes identified in GWAS for sleep/circadian rhythms and neurodevelopmental disorders. In the skyblue3 (3 genes) module (positively associated with LL), few hub-bottlenecks were identified (Fig. 9E). Lastly, the royal blue (3 genes) module (positively associated with poly IC) in females, also had few hub-bottlenecks, but some genes were homologous to human genes identified in GWAS for SCZ (*Stum*, *Lrrn2*), sleep (*Slc4A3, Rasd1)* and both SCZ and sleep (*Samd5, Hs6st3*) (Fig. 9F).

#### Distinct cell types are associated with different modules of co-expressed genes

We then aimed to determine the cell types involved in the gene co-expression networks identified through WGCNA. GSEA analyses on the LL DEGs lists yielded 1151 leading edge genes from the brown module, while GSEA analyses on the MIA DEGs lists yielded 959 and 210 leading edge genes from the brown and purple modules, respectively. LRcell analyses using these leading edge genes revealed that excitatory neurons of the subiculum (HC_2.7) and the dentate gyrus (HC_4.1 and 4.2) were driving the differential analysis of the brown module in LL (Supplementary Fig. 9A, while a neurogenesis neuron cell type enriched for *Sox4* and *Gabra5* (HC_13.5) was driving the differential expression of brown module genes in MIA (Supplementary Fig. 9B), and was also identified earlier using the full list of LL DEGs (Supplementary Fig. 7). In MIA, an excitatory neuronal cell type from the CA2/CA3 hippocampal sub-areas were implicated in the purple module (Supplementary Fig. 10).

As a complementary approach, we mapped the lists of hub-bottleneck genes from the brown, purple, green and cyan modules (as well as the whole lists of genes of the latter two modules, which contained fewer genes), on a published hippocampus/cortex single nucleus RNA sequencing dataset(55). Interestingly, some of these modules (brown, purple) included mainly hub-bottleneck genes primarily expressed in neurons (Supplementary Fig. 11, 12), whereas others (green, cyan) included genes (including hub-bottleneck genes) of non-neuronal cells (e.g. astrocytes, microglia, endothelial cells) (Supplementary Fig. 13, 14). These data point to the contribution of different cell types in the hippocampus in mediating the effects of MIA and LL.

## Discussion

Our study demonstrates that in mice prenatal MIA and circadian disruption during adolescence induce significant changes in behavior, microglial morphology, and transcriptome in a sex-dependent manner. Strikingly, LL exposure attenuated some of the effects of poly IC in behaviour (e.g. sociability), and hippocampal transcriptome. Large differences were apparent between sexes in behavior, microglia morphology, and transcriptome, with more effects in males than in females. WGCNA revealed significant gene modules positively associated with poly IC and LL in males, and fewer in females. Overall, this study shows that circadian disruption and abnormal light exposure can act as a potent modulator of the effects of a well-known risk factor of neurodevelopmental disorders, MIA.

Other studies similarly reported reduced anxiety-like behavior after LL (or dim light at night) in adult rodents (56–58), as well as reduced or no differences in corticosterone concentrations compared to 12:12LD(34, 57–59). Notably, LL did not lead to differences in anxiety-like behavior in poly IC-exposed mice. Reduced sociability was reported in some studies exploring prenatal poly IC(15), but not all(60). We found that poly IC-exposure led to reduced sociability compared to saline-exposed mice, but interestingly, poly IC and saline-exposed groups exposed to LL did not show these differences.

Previous work from our group found that in males, LL exposure during adulthood led to reduced sociability in poly IC exposed mice but had no effect on saline-exposed(35), whereas chronic jetlag exposure reduced sociability in both males and females (33). These data suggest an interplay between prenatal poly IC and LL exposure in males on both anxiety-like behavior and sociability. While cognitive impairments are a core feature of schizophrenia (61) and ASD(62), some individuals with ASD exhibit cognitive strengths, including attention and memory to details and a strong drive to detect patterns(63). Interestingly, there is an emerging literature on improved cognitive-like behavior in poly IC-exposed mice, whereby prenatal poly IC exposure conferred some functional advantages at relatively low doses(64). Thus, it is possible that this phenotype is present in our Barnes maze findings, where prenatal poly IC leads to better spatial memory, especially in the poly IC/LL group. Overall, in addition to being largely sex-specific, the effects of the factors were test-specific: in some cases, MIA and LL had opposite effects while in other cases the effects were additive. Also, when comparing with our previous study in which LL treatment was done in adulthood(35), here with LL treatment in adolescence, the effects of LL, and their interaction with MIA, were more prominent. This points to a neurodevelopmental effect of the lighting treatment.

Some individuals with schizophrenia and ASD exhibit characteristics of microglial activation and increased density in cortical regions (PFC, visual cortex), hippocampus and cerebellum(65). The current study showed that, in males, poly IC led to a more activated microglial profile and, additionally, while statistical significance was not reached, it appears that LL may have diminished these effects. A similar finding was also observed when poly IC and saline exposed mice were exposed to LL during adulthood(35). Interestingly, Hayes and colleagues found that MIA offspring had long-lived decreases in microglial immune reactivity across different developmental trajectories and replacing prenatal microglia could reverse the MIA-induced immune blunting(66). Additionally, diminished phagocytic function of microglia has been observed in response to 2 stressors that are independently known to stimulate microglia(67). Future studies would benefit from exploring beyond morphology and density and assess whether microglial functions are similarly diminished in poly IC exposed mice after LL exposure(68). Additional non-neuronal cell types in the hippocampus, such as the astrocytes, should also be investigated, as both MIA-and LL-associated modules (green, cyan) were found to include genes highly expressed not only in microglia but also astrocytes.

In the hippocampus transcriptome, more genes were found to be affected by prenatal poly IC in comparison to saline-exposure in males than in females, which is consistent with previous literature(60), and fewer DE genes were observed after LL exposure (in comparison to LD) in both sexes. WGCNA revealed that modules positively associated with LL exposure were enriched for different neuronal cell types, and for pathways involved in synaptic transmission and vesicle-mediated transport. Using the GWAS catalogue, not only were many of the central genes in these modules homologous to human genes related to sleep/circadian rhythms (e.g., *Prr7*) and neurodevelopmental disorders (e.g., *Erc2, Snca*), but they were also related to neurotransmission. For example, *Prr7* (*proline rich 7*) acts as a synapse-to-nucleus messenger to promote NMDA receptor-mediated excitotoxicity in neurons and, along with *Prr9*, is an essential component of circadian regulation of temperature(69). *Erc2* encodes a protein that belongs to the Rab3-interacting molecule (RIM)-binding protein family and is thought to be involved in the organization of the cytomatrix at the nerve terminals active zone, which regulates neurotransmitter release(70). Finally, *Snca (alpha-synuclein)* is a member of the synuclein family, it is found mainly in presynaptic terminals of neurons and plays a role in integrating presynaptic signaling and membrane trafficking(71). Thus, the mechanism by which LL affects behavior, microglia, and gene expression, could be related to disruptions in the expression of proteins required for efficient neurotransmitter release. Interestingly, the purple and brown WGCNA modules had an inverse association with MIA and LL, which is reminiscent of the opposite effects these factors had on some behaviors.

The multifactorial aspect of neurodevelopmental disorders, such as schizophrenia and ASD, prompts the study of the interaction between multiple risk factors. Previous work from our group found that mice prenatally exposed to poly IC (first ‘hit’ in utero) then subjected to LL (2^nd^ ‘hit’ in adulthood) show differences in behavior and microglia morphology (34). Building on these findings, the current study similarly employed the poly IC model, but this time the 2^nd^ ‘hit’ (LL exposure) was administered earlier during development (in adolescence). In humans, adolescence is a period when psychiatric disorders typically emerge (72), and it is marked by pronounced differences in the timing of sleep and circadian rhythms(36). Adolescence is also a time of heightened sensitivity to stressors. For example, exposure to social stress during adolescence led to more impairments in behavioral and neuronal outcomes than during adulthood(73–75). Thus, the impact of LL exposure, or at least its interaction with prenatal poly IC, appears to be influenced by age. Considering that adolescence is characterized by distinct changes in brain development, this finding could provide insight into specific pathways targeted by LL.

In this study, we investigated the interaction between MIA and circadian disruption. Our hypothesis was that environmental circadian disruption would act as a risk factor for NDDs and would exacerbate behavioral, cellular, and molecular impairments observed in the MIA mouse model. Though we provided evidence supporting a robust interaction between MIA and LL exposure, the relationship is evidently complex, and outcomes seem to vary depending on the specific measure being considered. Unexpectedly, for various measures, exposure to LL during adolescence resulted in a clear attenuation of the effects induced by MIA. While MIA is a well-established risk factor for NDDs, not all individuals exposed to MIA will develop these disorders. Susceptibility to neurodevelopmental disorders and the severity of symptoms are influenced by the complex interplay of various genetic and environmental factors. Our work demonstrates that in a mouse model of prenatal infection, disruptions in circadian rhythms induced by LL play a role in modulating the effects of MIA at behavioral, cellular, and molecular levels. Considering the pervasive exposure to environment circadian disruption imposed by modern society, investigating the impacts of such disruptions holds relevance for human populations. This is particularly the case for adolescents since their biological rhythms often shift toward later sleep/waking times, which may not align with daily routines, such as early school start times(36), this misalignment being further exacerbated by the use of electronic devices generating exposure of light at night(38, 39). Future studies may benefit from targeting specific mechanisms, as informed by data from mouse models, to gain a greater understanding on the impact of environmental circadian disruption and how it interacts with other risk factors. This knowledge could ultimately inform the development of circadian-based therapies.

## Acknowledgments

We thank the members of the Cermakian, Srivastava and Silveira laboratories for helpful discussions, Shashank Srikanta, Margaret Sayeh, Katherine Lord, Nicola Ludin and Joseph Rochford for help with behavioral tests or analysis, the Douglas animal care staff for tending to the animals, Melina Jaramillo Garcia and Bita Khadivjam at the Molecular and Cellular Microscopy Platform at the Douglas, and Elke Küster-Schöck at the Platform for Imaging by Microscopy at CHUSJ for help with cell analysis.

## Author contributions

Author contributions included conception and design of the experiments (TCD, LKS, PPS, NC), data acquisition (TCD, NH), data analysis (TCD, NO, DMA, DP, NH) and interpretation of results (TCD, NH, DMA, DP, NO, PPS, LKS, NC). Drafting manuscript and revising manuscript for publication (TCD, DMA, DP, NO, PPS, LKS, NC).

## Funding

This work was supported by grants from the Canadian Institutes of Health Research (PJT-153299 to NC and LKS) and Velux Stiftung (Project 927 to NC). McGill Ludmer-MI4 Collaborative Seed Fund Grant (to NC and PPS). TCD was supported by graduate scholarships from the Schizophrenia Society of Canada Foundation, the Canadian College of Neuropsychopharmacology, and the Fonds de Recherche du Québec – Santé. DP was supported by the HBHL Graduated Student Fellowship and NH was supported by an NSERC-CREATE undergraduate student research award.

## Competing Interests

The authors have nothing to disclose.

## Notes

### Competing Interest Statement

The authors have declared no competing interest.

## References

1. Jablensky A (2000): Epidemiology of schizophrenia: the global burden of disease and disability. Eur Arch Psychiatry Clin Neurosci. 250:274–285.

2. Baxter AJ, Brugha TS, Erskine HE, Scheurer RW, Vos T, Scott JG (2015): The epidemiology and global burden of autism spectrum disorders. Psychol Med. 45:601–613.

3. Fatemi SH, Folsom TD (2009): The neurodevelopmental hypothesis of schizophrenia, revisited. Schizophr Bull. 35:528–548.

4. Cheroni C, Caporale N, Testa G (2020): Autism spectrum disorder at the crossroad between genes and environment: contributions, convergences, and interactions in ASD developmental pathophysiology. Mol Autism. 11:69.

5. Schmitt A, Falkai P, Papiol S (2023): Neurodevelopmental disturbances in schizophrenia: evidence from genetic and environmental factors. Journal of Neural Transmission. 130:195–205.

6. Davis J, Eyre H, Jacka FN, Dodd S, Dean O, McEwen S, et al. (2016): A review of vulnerability and risks for schizophrenia: Beyond the two hit hypothesis. Neurosci Biobehav Rev. 65:185–194.

7. Kwon HK, Choi GB, Huh JR (2022): Maternal inflammation and its ramifications on fetal neurodevelopment. Trends Immunol. 43:230–244.

8. Massrali A, Adhya D, Srivastava DP, Baron-Cohen S, Kotter MR (2022): Virus-Induced Maternal Immune Activation as an Environmental Factor in the Etiology of Autism and Schizophrenia. Front Neurosci. 16:834058.

9. Brown AS, Begg MD, Gravenstein S, Schaefer CA, Wyatt RJ, Bresnahan M, et al. (2004): Serologic evidence of prenatal influenza in the etiology of schizophrenia. Arch Gen Psychiatry. 61:774–780.

10. Lee BK, Magnusson C, Gardner RM, Blomstrom A, Newschaffer CJ, Burstyn I, et al. (2015): Maternal hospitalization with infection during pregnancy and risk of autism spectrum disorders. Brain Behav Immun. 44:100–105.

11. Rapoport JL, Giedd JN, Gogtay N (2012): Neurodevelopmental model of schizophrenia: update 2012. Mol Psychiatry. 17:1228–1238.

12. Haddad FL, Patel SV, Schmid S (2020): Maternal Immune Activation by Poly I:C as a preclinical Model for Neurodevelopmental Disorders: A focus on Autism and Schizophrenia. Neuroscience & Biobehavioral Reviews. 113:546–567.

13. Meyer U (2014): Prenatal poly(i:C) exposure and other developmental immune activation models in rodent systems. Biol Psychiatry. 75:307–315.

14. Murray BG, Davies DA, Molder JJ, Howland JG (2017): Maternal immune activation during pregnancy in rats impairs working memory capacity of the offspring. Neurobiol Learn Mem. 141:150–156.

15. Hui CW, St-Pierre A, El Hajj H, Remy Y, Hebert SS, Luheshi GN, et al. (2018): Prenatal Immune Challenge in Mice Leads to Partly Sex-Dependent Behavioral, Microglial, and Molecular Abnormalities Associated with Schizophrenia. Front Mol Neurosci. 11:13.

16. Paolicelli RC, Ferretti MT (2017): Function and Dysfunction of Microglia during Brain Development: Consequences for Synapses and Neural Circuits. Front Synaptic Neurosci. 9:9.

17. Knuesel I, Chicha L, Britschgi M, Schobel SA, Bodmer M, Hellings JA, et al. (2014): Maternal immune activation and abnormal brain development across CNS disorders. Nat Rev Neurol. 10:643–660.

18. Giovanoli S, Notter T, Richetto J, Labouesse MA, Vuillermot S, Riva MA, et al. (2015): Late prenatal immune activation causes hippocampal deficits in the absence of persistent inflammation across aging. J Neuroinflammation. 12:221.

19. Schaafsma W, Basterra LB, Jacobs S, Brouwer N, Meerlo P, Schaafsma A, et al. (2017): Maternal inflammation induces immune activation of fetal microglia and leads to disrupted microglia immune responses, behavior, and learning performance in adulthood. Neurobiol Dis. 106:291–300.

20. Patke A, Young MW, Axelrod S (2020): Molecular mechanisms and physiological importance of circadian rhythms. Nat Rev Mol Cell Biol. 21:67–84.

21. Hastings MH, Maywood ES, Brancaccio M (2018): Generation of circadian rhythms in the suprachiasmatic nucleus. Nat Rev Neurosci. 19:453–469.

22. Karatsoreos IN, Bhagat S, Bloss EB, Morrison JH, McEwen BS (2011): Disruption of circadian clocks has ramifications for metabolism, brain, and behavior. Proc Natl Acad Sci U S A. 108:1657–1662.

23. Walker WH, 2nd, Walton JC, DeVries AC, Nelson RJ (2020): Circadian rhythm disruption and mental health. Transl Psychiatry. 10:28.

24. Krakowiak P, Goodlin-Jones B, Hertz-Picciotto I, Croen LA, Hansen RL (2008): Sleep problems in children with autism spectrum disorders, developmental delays, and typical development: a population-based study. J Sleep Res. 17:197–206.

25. Wulff K, Dijk DJ, Middleton B, Foster RG, Joyce EM (2012): Sleep and circadian rhythm disruption in schizophrenia. Br J Psychiatry. 200:308–316.

26. Johansson AS, Owe-Larsson B, Hetta J, Lundkvist GB (2016): Altered circadian clock gene expression in patients with schizophrenia. Schizophr Res. 174:17–23.

27. Seney ML, Cahill K, Enwright JF, 3rd, Logan RW, Huo Z, Zong W, et al. (2019): Diurnal rhythms in gene expression in the prefrontal cortex in schizophrenia. Nat Commun. 10:3355.

28. Delorme TC, Srivastava LK, Cermakian N (2020): Are Circadian Disturbances a Core Pathophysiological Component of Schizophrenia? J Biol Rhythms. 35:325–339.

29. Wintler T, Schoch H, Frank MG, Peixoto L (2020): Sleep, brain development, and autism spectrum disorders: Insights from animal models. J Neurosci Res. 98:1137–1149.

30. Delorme TC, Srivastava LK, Cermakian N (2021): Altered circadian rhythms in a mouse model of neurodevelopmental disorders based on prenatal maternal immune activation. Brain Behav Immun. 93:119–131.

31. Tandon R, Gaebel W, Barch DM, Bustillo J, Gur RE, Heckers S, et al. (2013): Definition and description of schizophrenia in the DSM-5. Schizophr Res. 150:3–10.

32. Boivin DB (2000): Influence of sleep-wake and circadian rhythm disturbances in psychiatric disorders. J Psychiatry Neurosci. 25:446–458.

33. Cloutier ME, Srivastava LK, Cermakian N (2022): Exposure to Circadian Disruption During Adolescence Interacts With a Genetic Risk Factor to Modify Schizophrenia-relevant Behaviors in a Sex-dependent Manner. J Biol Rhythms. 37:655–672.

34. Bhardwaj SK, Stojkovic K, Kiessling S, Srivastava LK, Cermakian N (2015): Constant light uncovers behavioral effects of a mutation in the schizophrenia risk gene Dtnbp1 in mice. Behav Brain Res. 284:58–68.

35. Delorme TC, Ozell-Landry W, Cermakian N, Srivastava LK (2023): Behavioral and cellular responses to circadian disruption and prenatal immune activation in mice. Scientific Reports. 13:7791.

36. Logan RW, McClung CA (2019): Rhythms of life: circadian disruption and brain disorders across the lifespan. Nat Rev Neurosci. 20:49–65.

37. Perkinson-Gloor N, Lemola S, Grob A (2013): Sleep duration, positive attitude toward life, and academic achievement: The role of daytime tiredness, behavioral persistence, and school start times. Journal of Adolescence. 36:311–318.

38. Fonken LK, Nelson RJ (2016): Effects of light exposure at night during development. Current Opinion in Behavioral Sciences. 7:33–39.

39. Spanagel R (2018): Alterations of the Biological Clock May Contribute to the Emergence of Mental Disorders During Adolescence. Biol Psychiatry. 83:978–980.

40. Ochoa S, Usall J, Cobo J, Labad X, Kulkarni J (2012): Gender Differences in Schizophrenia and First-Episode Psychosis: A Comprehensive Literature Review. Schizophrenia Research and Treatment. 2012:1–9.

41. Ferri SL, Abel T, Brodkin ES (2018): Sex Differences in Autism Spectrum Disorder: a Review. Curr Psychiatry Rep. 20:9.

42. Xuan IC, Hampson DR (2014): Gender-dependent effects of maternal immune activation on the behavior of mouse offspring. PLoS One. 9:e104433.

43. Valle FP (1970): Effects of strain, sex, and illumination on open-field behavior of rats. Am J Psychol. 83:103–111.

44. Kaidanovich-Beilin O, Lipina T, Vukobradovic I, Roder J, Woodgett JR (2011): Assessment of Social Interaction Behaviors. Journal of Visualized Experiments.e2473.

45. Paxinos G, Franklin KB (2019): Paxinos and Franklin’s the mouse brain in stereotaxic coordinates. Academic press.

46. Love MI, Huber W, Anders S (2014): Moderated estimation of fold change and dispersion for RNA-seq data with DESeq2. Genome Biology. 15:550.

47. Cahill KM, Huo Z, Tseng GC, Logan RW, Seney ML (2018): Improved identification of concordant and discordant gene expression signatures using an updated rank-rank hypergeometric overlap approach. Scientific Reports. 8:9588.

48. Langfelder P, Horvath S (2008): WGCNA: an R package for weighted correlation network analysis. BMC bioinformatics. 9:1–13.

49. de Lima RMS, Barth B, Mar Arcego D, de Mendonça Filho EJ, Patel S, Wang Z, et al. (2022): Leptin receptor co-expression gene network moderates the effect of early life adversity on eating behavior in children. Communications Biology. 5:1092.

50. Yu H, Kim PM, Sprecher E, Trifonov V, Gerstein M (2007): The Importance of Bottlenecks in Protein Networks: Correlation with Gene Essentiality and Expression Dynamics. PLoS Computational Biology. 3:e59.

51. Neves De Oliveira B-H, Dalmaz C, Zeidán-Chuliá F (2018): Network-Based Identification of Altered Stem Cell Pluripotency and Calcium Signaling Pathways in Metastatic Melanoma. Medical Sciences. 6:23.

52. Ma W, Sharma S, Jin P, Gourley SL, Qin ZS (2022): LRcell: detecting the source of differential expression at the sub-cell-type level from bulk RNA-seq data. Brief Bioinform. 23.

53. Saunders A, Macosko EZ, Wysoker A, Goldman M, Krienen FM, de Rivera H, et al. (2018): Molecular Diversity and Specializations among the Cells of the Adult Mouse Brain. Cell. 174:1015–1030.e1016.

54. Program CS-CB, Abdulla S, Aevermann B, Assis P, Badajoz S, Bell SM, et al. (2023): CZ CELL×GENE Discover: A single-cell data platform for scalable exploration, analysis and modeling of aggregated data. bioRxiv. 2023.2010.2030.563174.

55. Yao Z, van Velthoven CTJ, Nguyen TN, Goldy J, Sedeno-Cortes AE, Baftizadeh F, et al. (2021): A taxonomy of transcriptomic cell types across the isocortex and hippocampal formation. Cell. 184:3222–3241.e3226.

56. Ma W-P, Cao J, Tian M, Cui M-H, Han H-L, Yang Y-X, et al. (2007): Exposure to chronic constant light impairs spatial memory and influences long-term depression in rats. Neuroscience Research. 59:224–230.

57. Fonken LK, Finy MS, Walton JC, Weil ZM, Workman JL, Ross J, et al. (2009): Influence of light at night on murine anxiety- and depressive-like responses. Behav Brain Res. 205:349–354.

58. Bedrosian TA, Fonken LK, Walton JC, Haim A, Nelson RJ (2011): Dim light at night provokes depression-like behaviors and reduces CA1 dendritic spine density in female hamsters. Psychoneuroendocrinology. 36:1062–1069.

59. Claustrat B, Valatx JL, Harthé C, Brun J (2008): Effect of Constant Light on Prolactin and Corticosterone Rhythms Evaluated Using a Noninvasive Urine Sampling Protocol in the Rat. Hormone and Metabolic Research. 40:398–403.

60. Guma E, Bordignon PDC, Devenyi GA, Gallino D, Anastassiadis C, Cvetkovska V, et al. (2021): Early or Late Gestational Exposure to Maternal Immune Activation Alters Neurodevelopmental Trajectories in Mice: An Integrated Neuroimaging, Behavioral, and Transcriptional Study. Biological Psychiatry. 90:328–341.

61. McCutcheon RA, Keefe RSE, McGuire PK (2023): Cognitive impairment in schizophrenia: aetiology, pathophysiology, and treatment. Molecular Psychiatry. 28:1902–1918.

62. Banker SM, Gu X, Schiller D, Foss-Feig JH (2021): Hippocampal contributions to social and cognitive deficits in autism spectrum disorder. Trends Neurosci. 44:793–807.

63. Baron-Cohen S (2017): Editorial Perspective: Neurodiversity - a revolutionary concept for autism and psychiatry. J Child Psychol Psychiatry. 58:744–747.

64. Zhao X, Tran H, Derosa H, Roderick RC, Kentner AC (2021): Hidden talents: Poly (I:C)Linduced maternal immune activation improves mouse visual discrimination performance and reversal learning in a sexLdependent manner. *Genes*, Brain and Behavior. 20:e12755.

65. Bloomfield PS, Selvaraj S, Veronese M, Rizzo G, Bertoldo A, Owen DR, et al. (2016): Microglial Activity in People at Ultra High Risk of Psychosis and in Schizophrenia: An [11C] PBR28 PET Brain Imaging Study. Am J Psychiatry. 173:44–52.

66. Hayes LN, An K, Carloni E, Li F, Vincent E, Trippaers C, et al. (2022): Prenatal immune stress blunts microglia reactivity, impairing neurocircuitry. Nature. 610:327–334.

67. Block CL, Eroglu O, Mague SD, Smith CJ, Ceasrine AM, Sriworarat C, et al. (2022): Prenatal environmental stressors impair postnatal microglia function and adult behavior in males. Cell Reports. 40:111161.

68. Paolicelli RC, Sierra A, Stevens B, Tremblay ME, Aguzzi A, Ajami B, et al. (2022): Microglia states and nomenclature: A field at its crossroads. Neuron. 110:3458–3483.

69. Kravchick DO, Karpova A, Hrdinka M, LopezLRojas J, Iacobas S, Carbonell AU, et al. (2016): Synaptonuclear messenger <scp>PRR</scp> 7 inhibits cLJun ubiquitination and regulates <scp>NMDA</scp> Lmediated excitotoxicity. The EMBO Journal. 35:1923–1934.

70. Wang Y, Liu X, Biederer T, Südhof TC (2002): A family of RIM-binding proteins regulated by alternative splicing: Implications for the genesis of synaptic active zones. Proceedings of the National Academy of Sciences. 99:14464–14469.

71. Bernal-Conde LD, Ramos-Acevedo R, Reyes-Hernández MA, Balbuena-Olvera AJ, Morales-Moreno ID, Argüero-Sánchez R, et al. (2020): Alpha-Synuclein Physiology and Pathology: A Perspective on Cellular Structures and Organelles. Frontiers in Neuroscience. 13:1399.

72. Kessler RC, Amminger GP, Aguilar-Gaxiola S, Alonso J, Lee S, Ustun TB (2007): Age of onset of mental disorders: a review of recent literature. Current Opinion in Psychiatry. 20:359–364.

73. Morrissey MD, Mathews IZ, McCormick CM (2011): Enduring deficits in contextual and auditory fear conditioning after adolescent, not adult, social instability stress in male rats. Neurobiology of Learning and Memory. 95:46–56.

74. McCormick CM, Mongillo DL, Simone JJ (2013): Age and adolescent social stress effects on fear extinction in female rats. Stress. 16:678–688.

75. Ver Hoeve ES, Kelly G, Luz S, Ghanshani S, Bhatnagar S (2013): Short-term and long-term effects of repeated social defeat during adolescence or adulthood in female rats. Neuroscience. 249:63–73.

