## Supplemental Material for "Effects of prenatal maternal immune activation and exposure to circadian disruption during adolescence: exploring the two-hit model of neurodevelopmental disorders"

### Supplementary Material

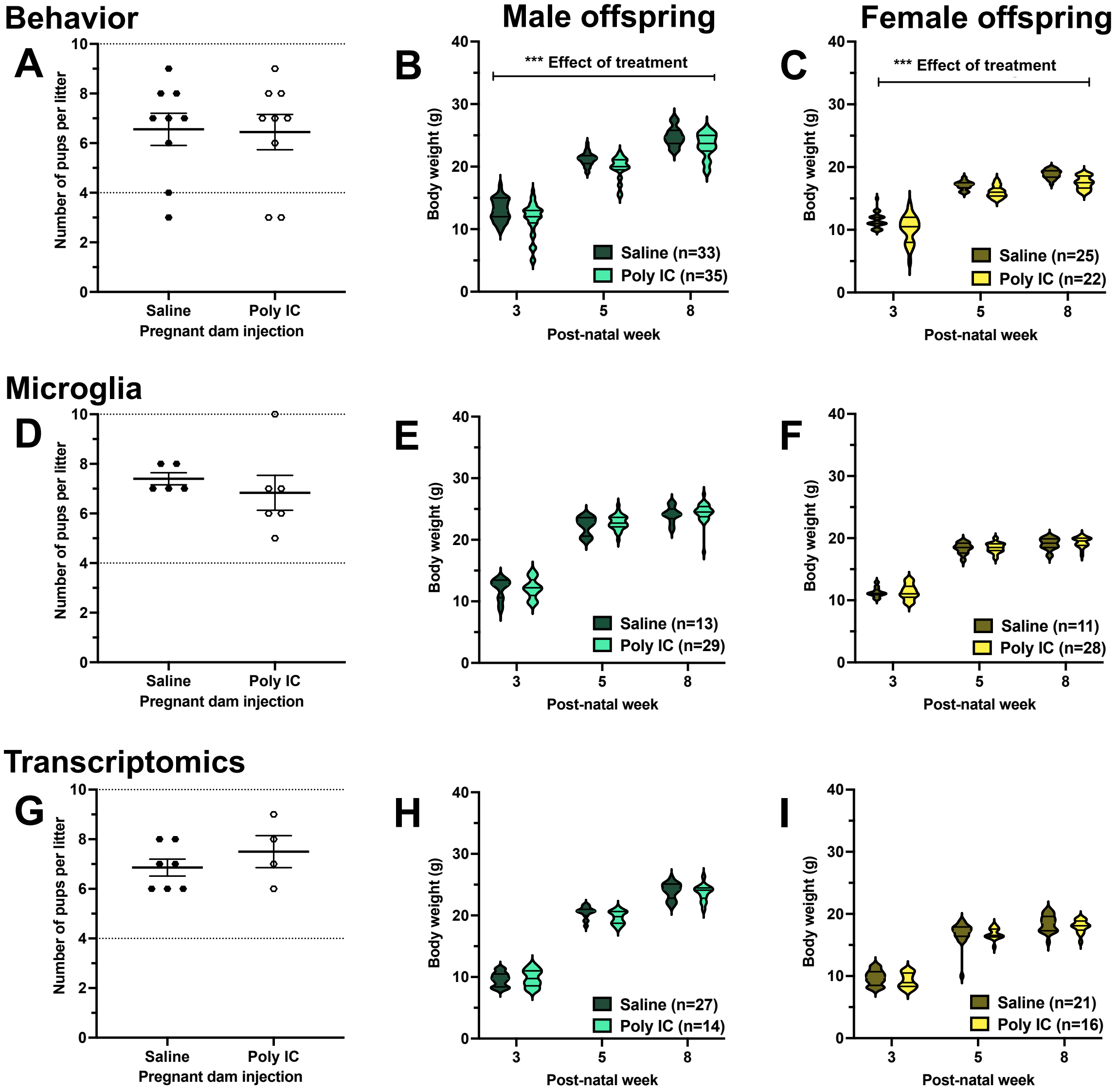


**Supplementary Fig. 1.** Litters of poly IC- and saline-exposed dams did not differ in number of pups for the litters used for behavioral testing (**A**), microglia analysis **(D)** or transcriptomics **(G).** Body weight was assessed for each cohort used for behavioral testing (**B, C**), microglia analysis **(E,F)** or transcriptomics **(H, I)** in male **(B, E, H)** and female **(C, F, I)** offspring at 3, 5, and 8 post-natal weeks. Two-way ANOVAs, with treatment (poly IC, saline) as a between-subjects variable and age (post-natal week 3, 5, 8) as a within-subjects variable were conducted. Treatment x age interactions and main effects for each sex separately were assessed.

#### Poly IC validation

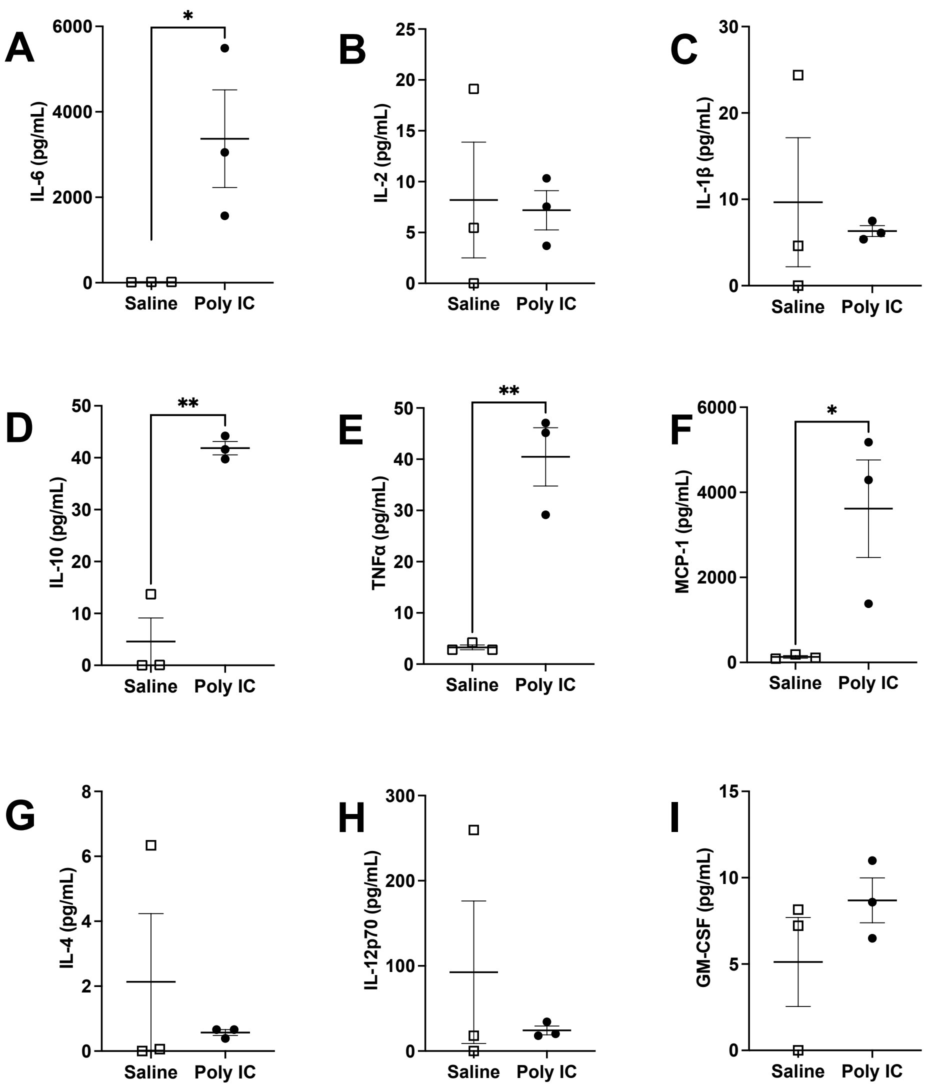


**Supplementary Fig. 2. Pro- and anti-inflammatory cytokine and chemokine expression levels in plasma of pregnant dams injected with poly IC (n = 3) or saline (n = 3).** Expression levels of the following cytokine and chemokines: (**A**) IL-6, (**B**) IL-2, (**C**) IL-1β, (**D**)IL-10, (**E**) TNFα (**F**) MCP-1, (**G**) IL-4 (**H**) IL-12(p70) and (**I**) GM-CSF. Samples that were below the detectable range were designated as 0pg/mL. Individual data points represent independent dams and data were represented as mean ± SEM. An independent samples t-test (with Welsh's correction, or Mann-Whitney U test when appropriate) was used, $**p < 0.01$, and $*p < 0.05$.

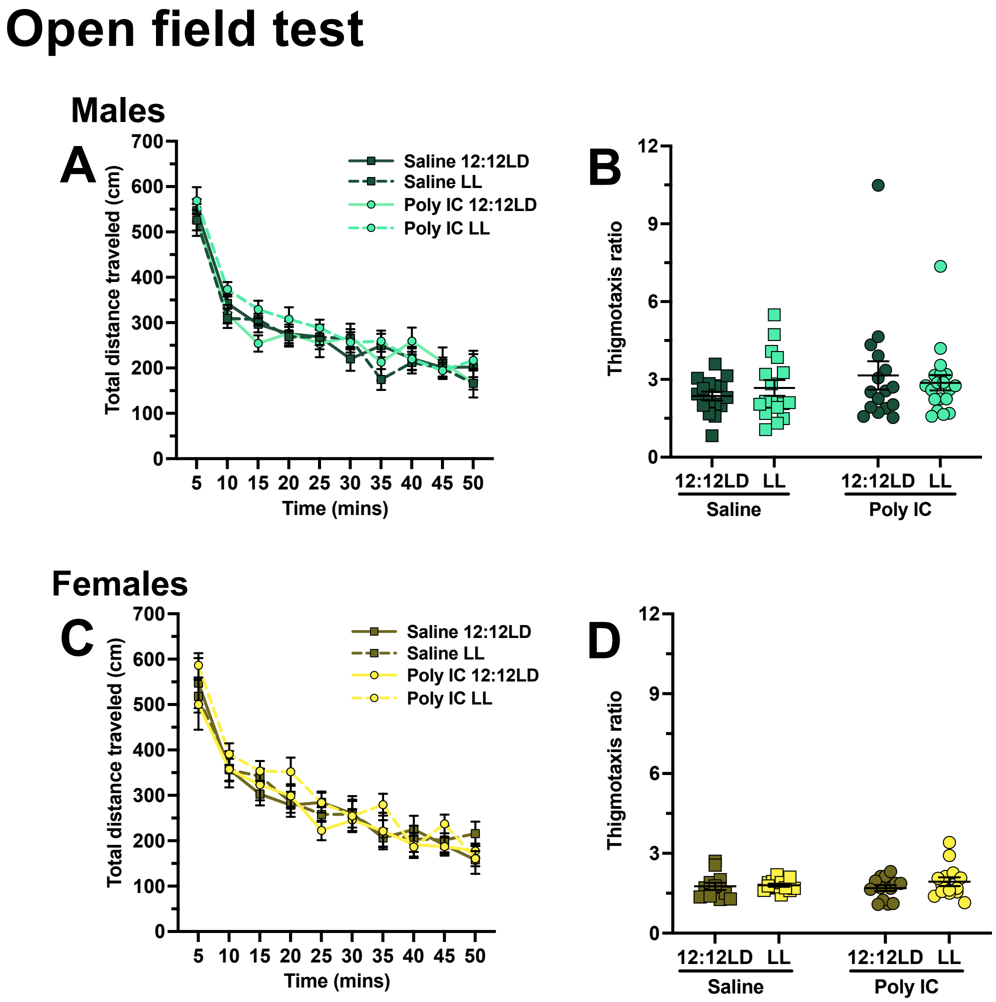


**Supplementary Fig. 3. Similar spontaneous locomotion in saline and poly IC-exposed mice.** Thigmotaxis **(A, C)** and total distance traveled (**B, D**) were assessed in males (**A-B**) and females (**C-D**). For panels **A and C** data points represent individual mice and are presented as mean ± SEM. Two-way ANOVAs (factors treatment x lighting with Tukey's post-hoc comparisons) were conducted. For panels **B** and **D,** group averages ± SEM are shown over each 5-minute bin of the test. Three-way ANOVAs (factors treatment x lighting x time) were conducted.

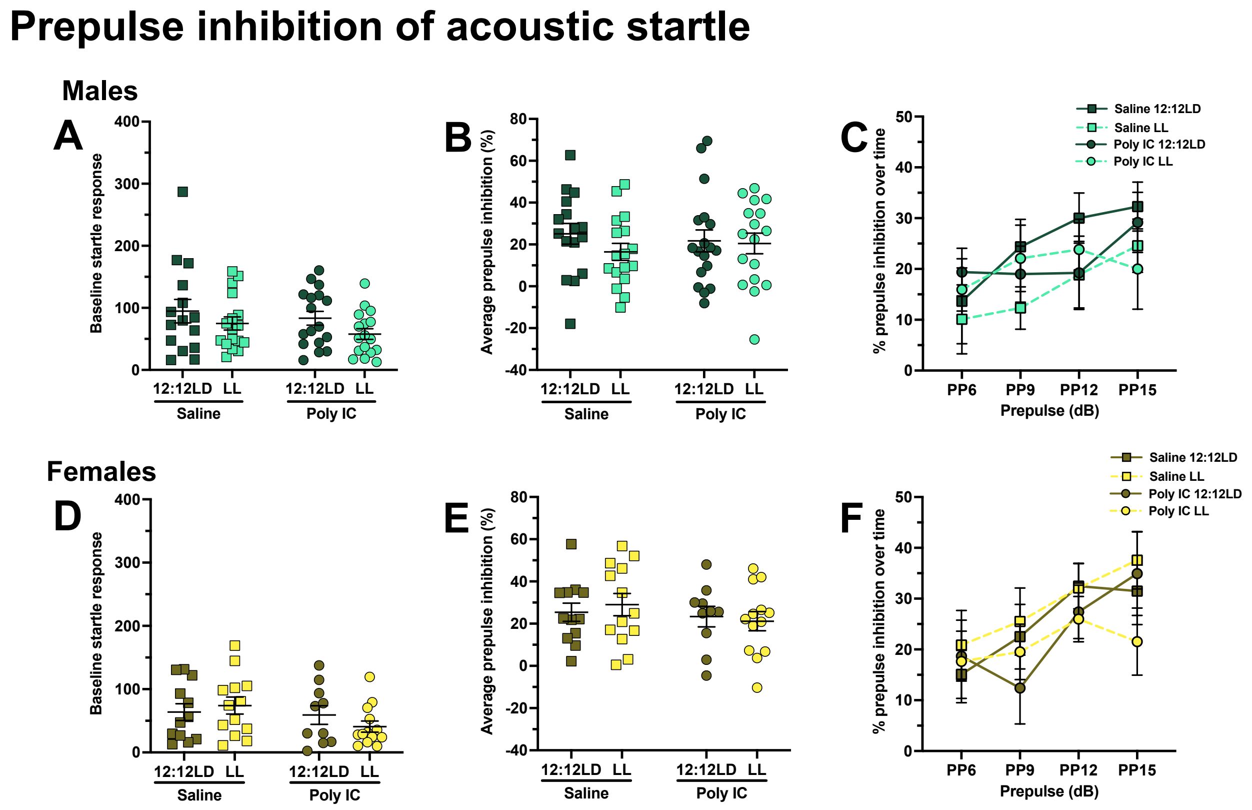


**Supplementary Fig. 4. No prepulse inhibition of acoustic startle (PPI) deficits following poly IC exposure.** Baseline startle response (**A, D**), average PPI (%) (**B, E**) and PPI (%) across each prepulse levels (**C, F**) were assessed in males (**A-C**) and females (**D-F**). For panels **A, B, D and E** data points represent individual mice and are presented as mean ± SEM. Two-way ANOVAs (factors treatment x lighting with Tukey's post-hoc comparisons) were conducted. For panels **C** and **F,** group averages ± SEM are shown over prepulse level. Three-way ANOVAs (factors treatment x lighting x time) were conducted.

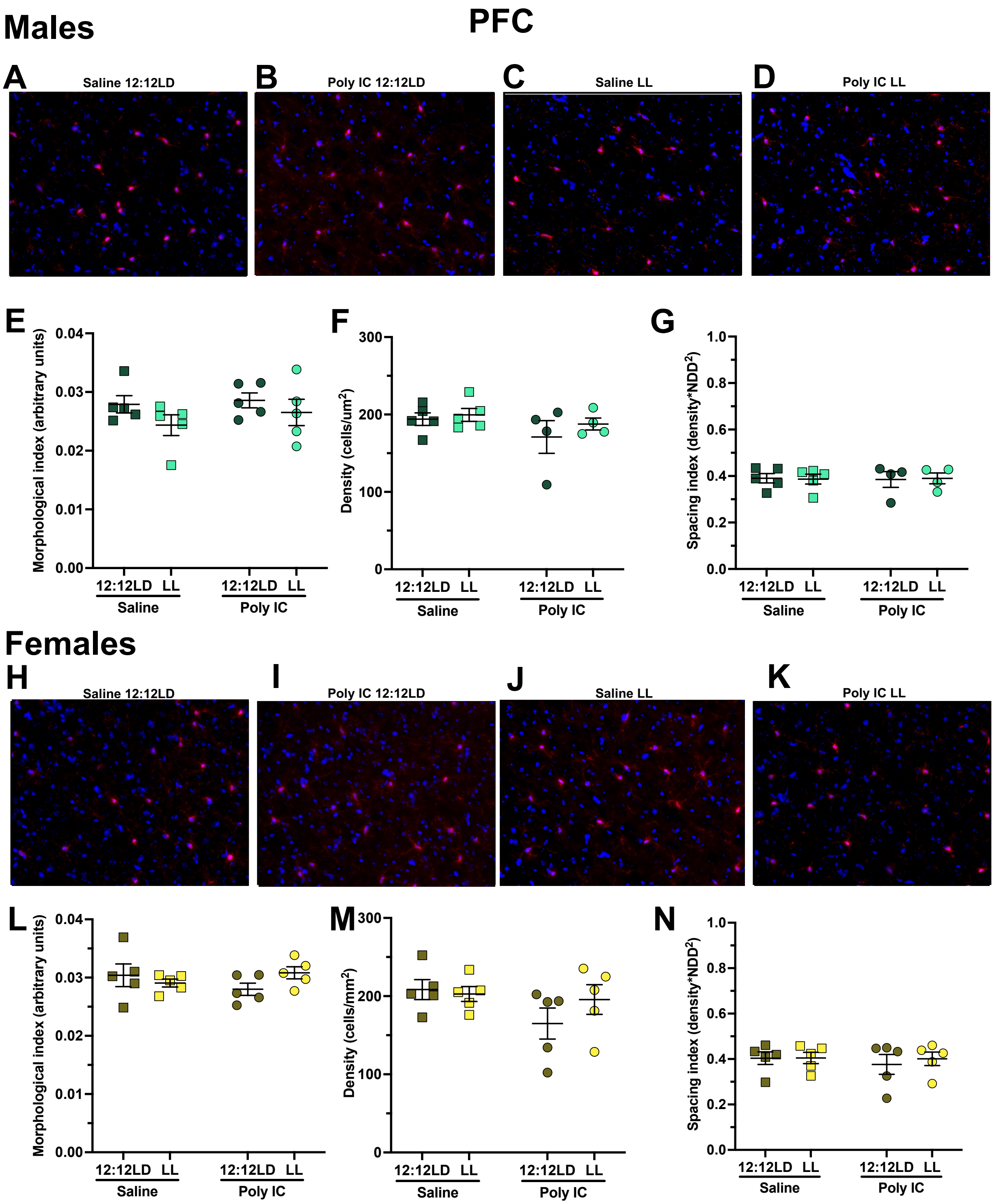


**Supplementary Fig. 5. Poly IC exposure did not lead to differences in morphological index or density in the prefrontal cortex.** Representative images of microglia from the DG for each group under each lighting condition are shown for males **(A-D)** and females **(H-K)**. Morphological index (**E, L**), density (**F, M**) and spacing index (**G, N**) were assessed for males (**E-G**) and females (**L-N**). Data points represent individual mice and are presented as mean ± SEM. Two-way ANOVAs (factors treatment x lighting with Tukey's post-hoc comparisons) were conducted.

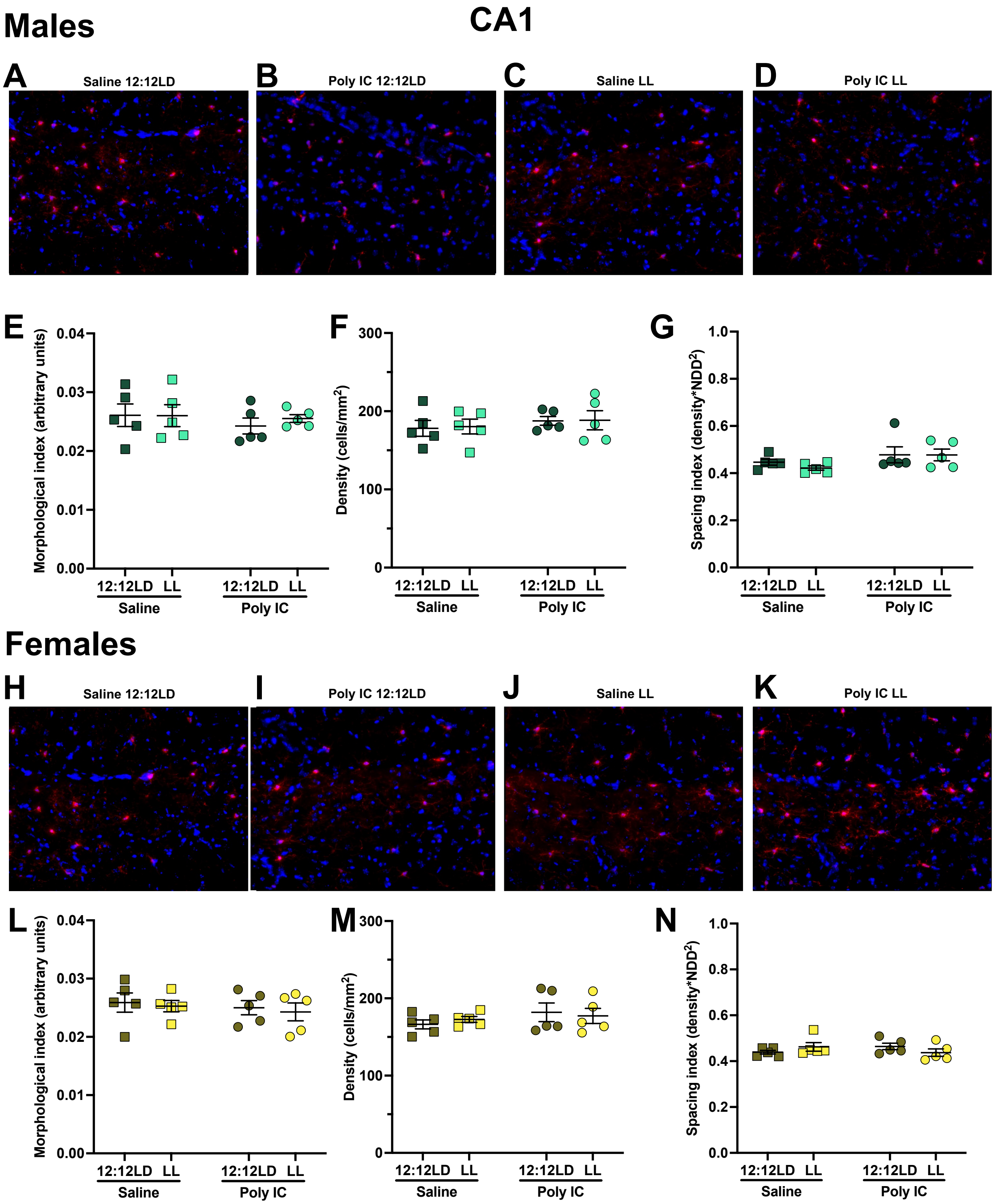


**Supplementary Fig. 6. Poly IC exposure did not lead to differences in morphological index or density in CA1.** Representative images of microglia from the DG for each group under each lighting condition are shown for males **(A-D)** and females **(H-K)**. Morphological index (**E, L**), density (**F, M**) and spacing index (**G, N**) were assessed for males (**E-G**) and females (**L-N**). Data points represent individual mice and are presented as mean ± SEM. Two-way ANOVAs (factors treatment x lighting with Tukey's post-hoc comparisons) were conducted.

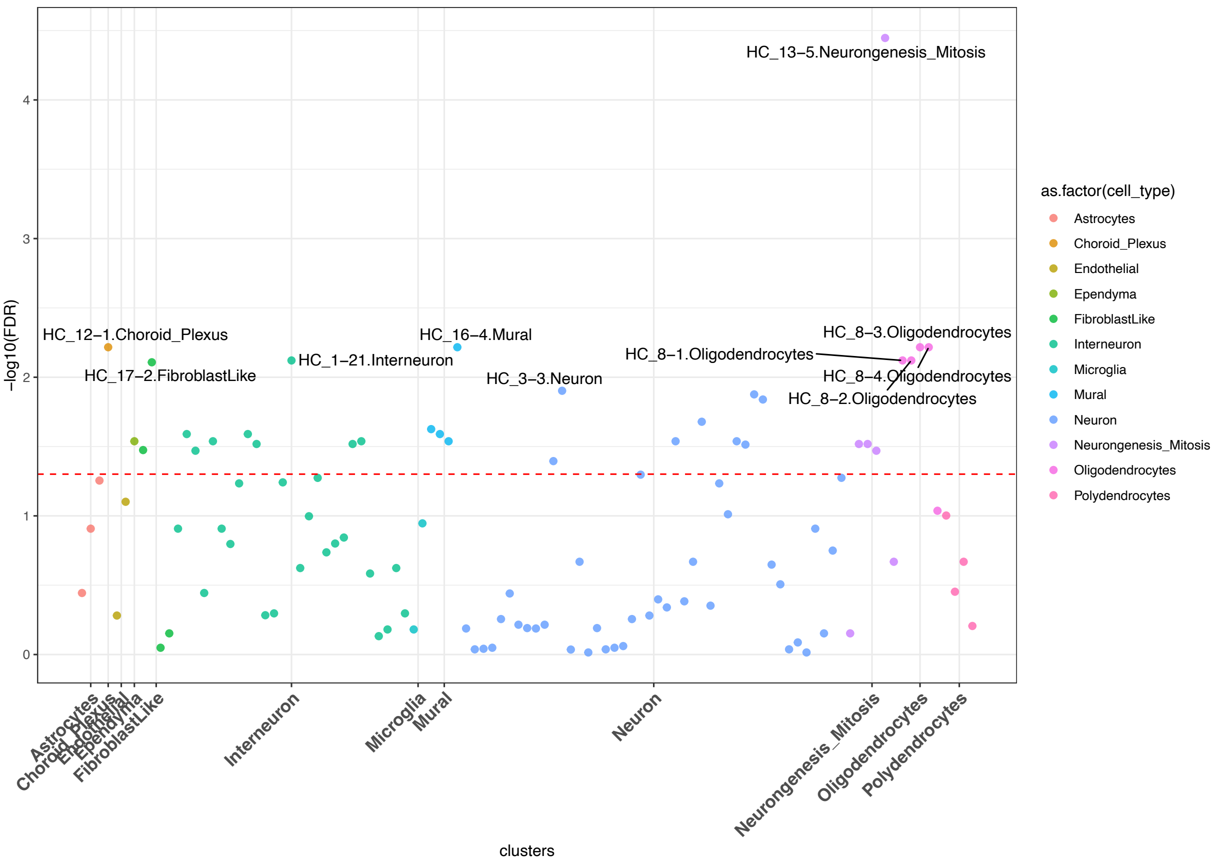


**Supplementary Figure 7. Manhattan plot of the mouse hippocampal cell types driving the differential expression in LL.** The entire list of differentially expressed genes between controls and LD and LL was used. The dotted red line represents the significant cut-off, and each dot represent different cell subtypes.

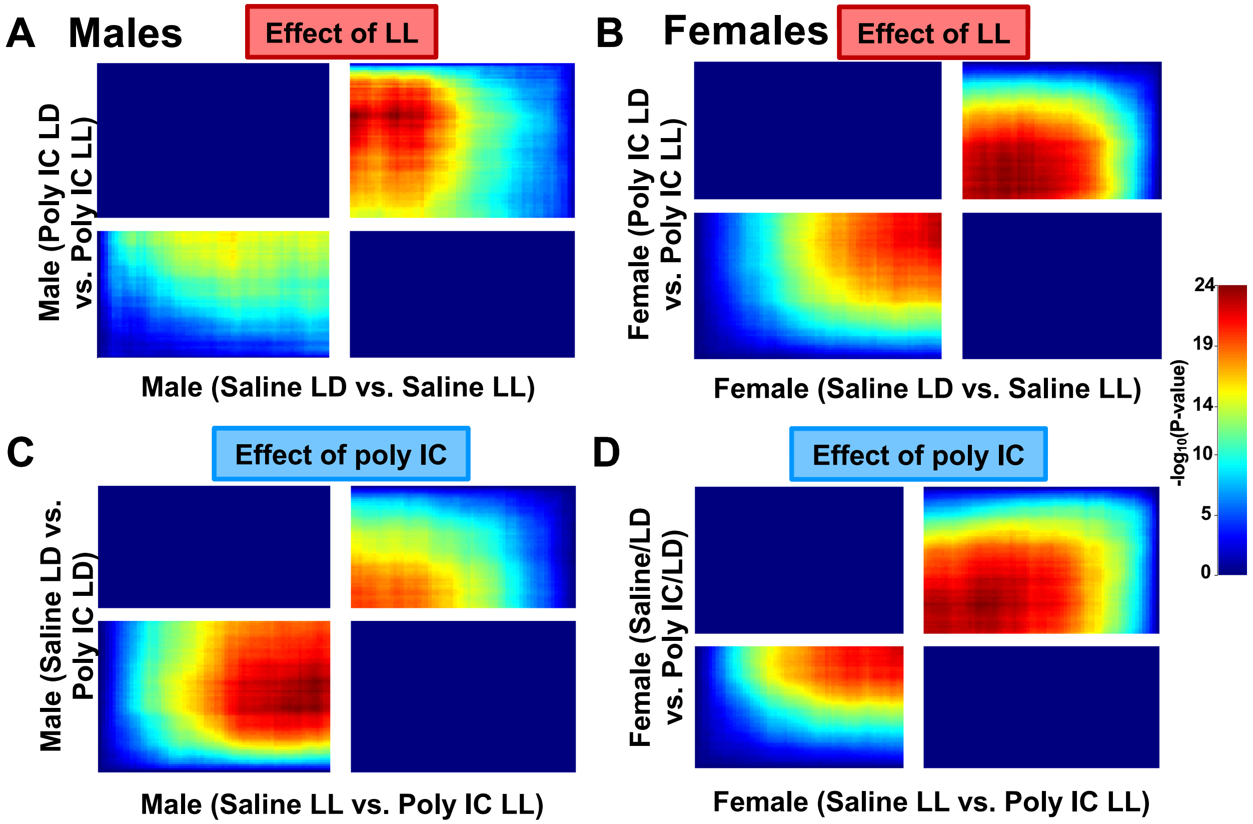


**Supplementary Fig. 8. Rank rank hypergeometric overlap 2 (RRHO2) analysis results in concordant patterns between treatment conditions for each sex.**

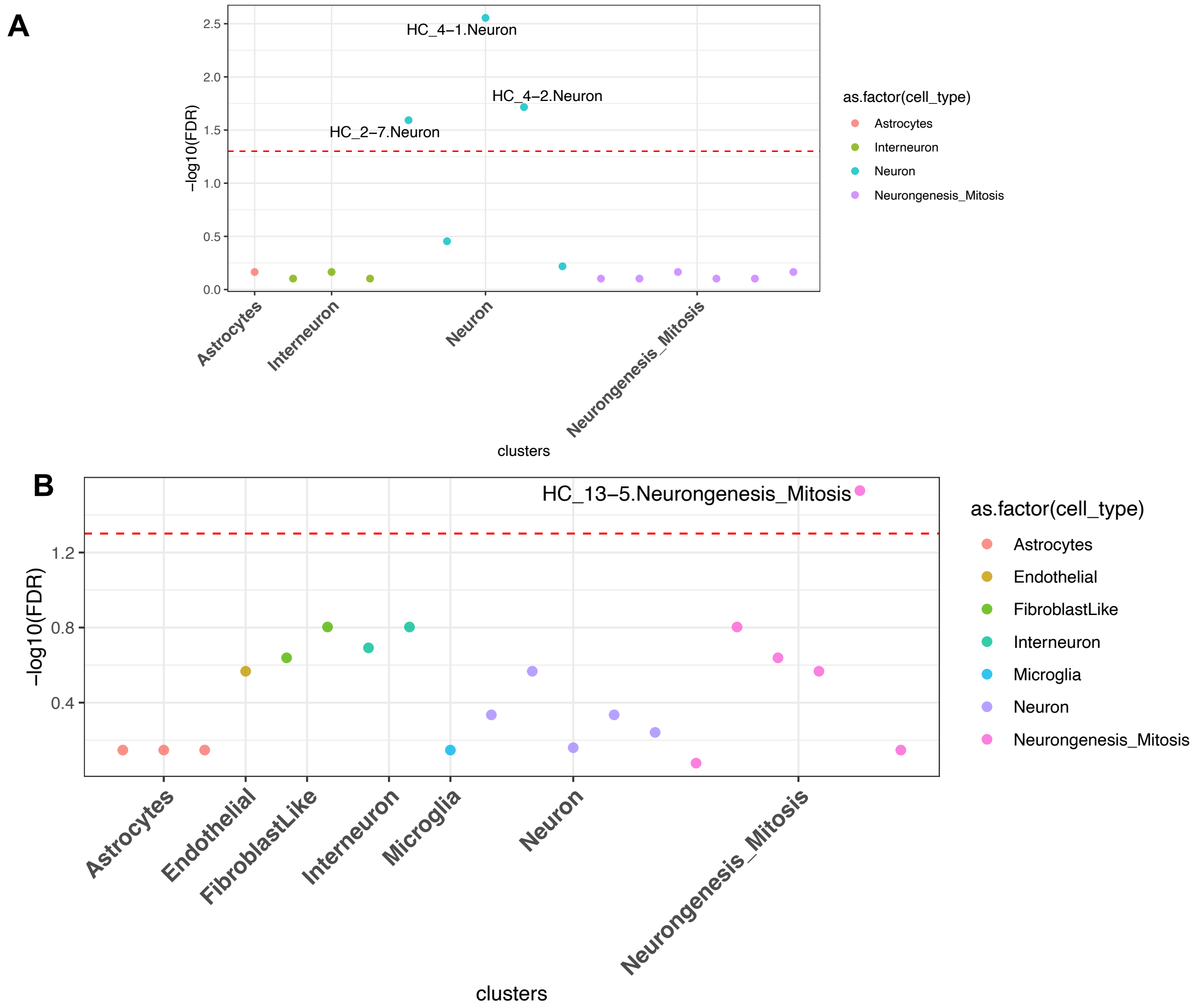


**Supplementary Figure 9. Manhattan plot of the mouse hippocampal cell types driving the differential expression of genes from the brown module.** The brown module genes chosen were enriched in among the differentially expressed genes in LL (A) and MIA (B). The dotted red line represents the significant cut-off, and each dot represent different cell subtypes.

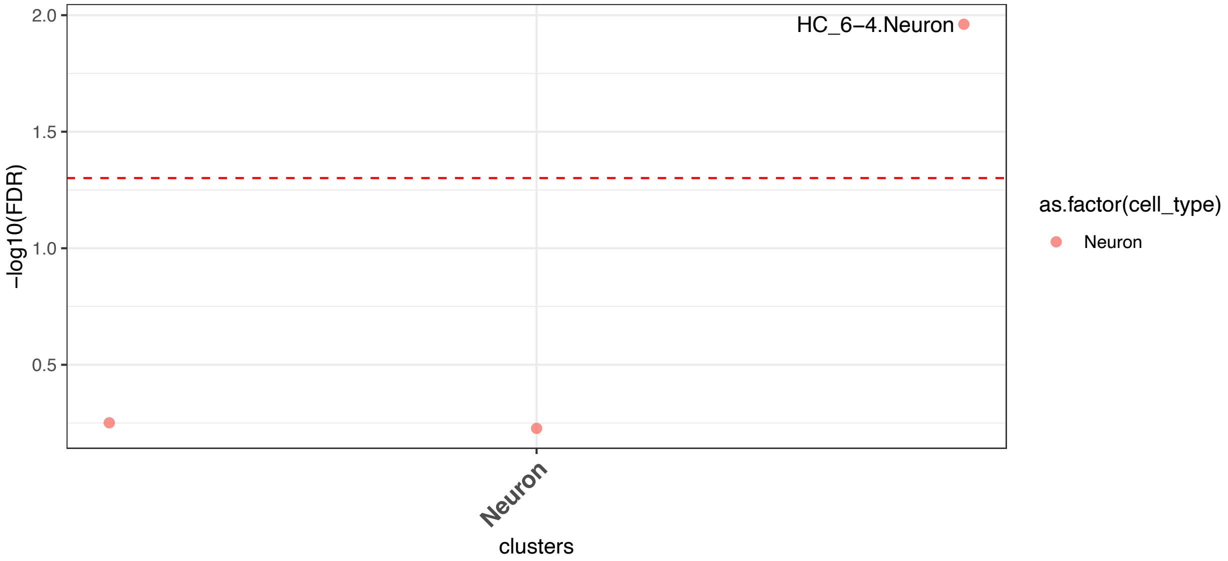


**Supplementary Figure 10. Manhattan plot of the mouse hippocampal cell types driving the differential expression of genes from the purple module in MIA.** The dotted red line represents the significant cut-off, and each dot represent different cell subtypes.

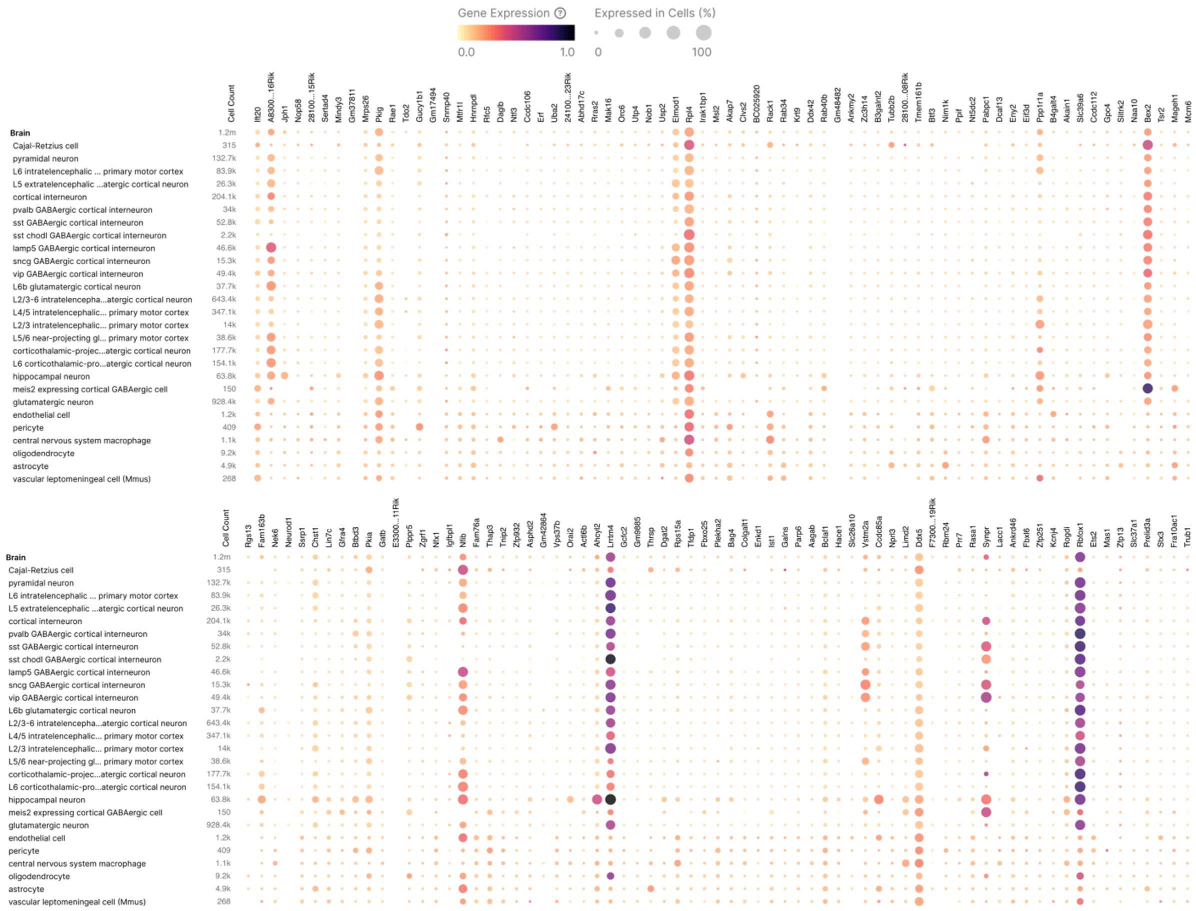

**Supplementary Figure 11. Dot plot representation of the gene expression of the brown module hub-bottleneck genes across cell types in the mouse hippocampus and isocortex.**

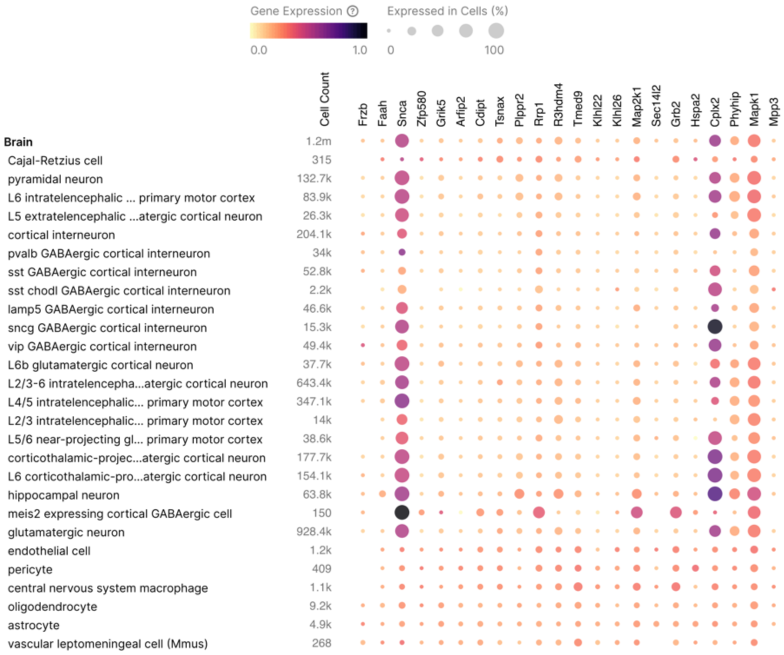


**Supplementary Figure 12. Dot plot representation of the gene expression of the purple module hub-bottleneck genes across cell types in the mouse hippocampus and isocortex.**

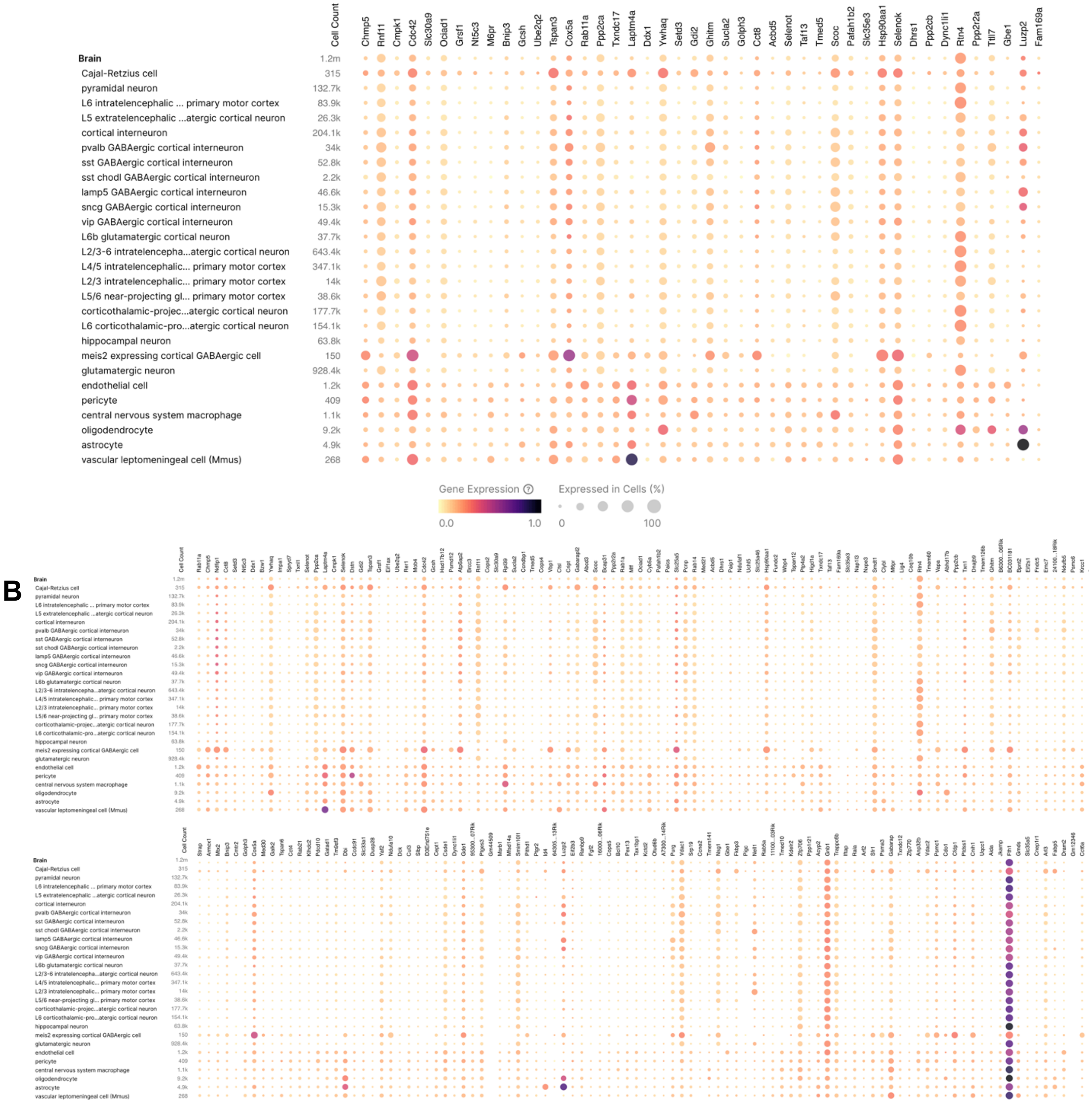


**Supplementary Figure 13. Dot plot representation of the gene expression of the green module genes across cell types in the mouse hippocampus and isocortex.** The dot plots were created with the hub-bottleneck genes only (A) and all genes (B) in the green module.

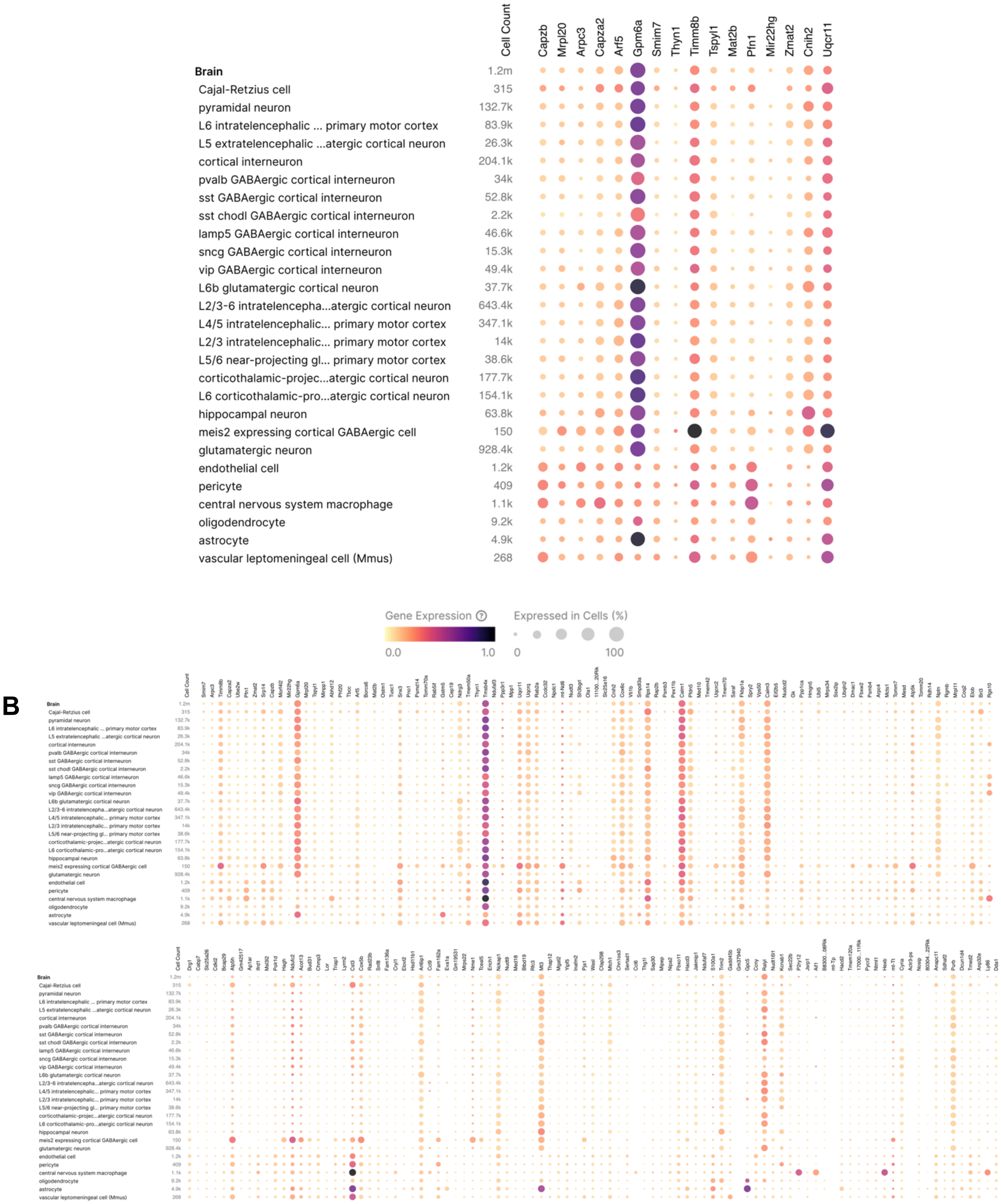

**Supplementary Figure 14. Dot plot representation of the gene expression of the cyan module genes across cell types in the mouse hippocampus and isocortex.** The dot plots were created with the hub-bottleneck genes only (A) and all genes (B) in the cyan module.

**Supplementary Table 1. Sex differences in behavior of poly IC and saline-exposed mice under LD or LL.**

| | Treatment x Sex x Lighting | | Treatment x Sex | | Lighting x Sex | | Main Effect Sex | |
|---|---|---|---|---|---|---|---|---|
| | F | *p-value* | F | *p-value* | F | *p-value* | F | *p-value* |
| Open field | | | | | | | | |
| Thigmotaxis ratio | 0.9472 | 0.3325 | 1.2780 | 0.2606 | 0.0971 | 0.7558 | 22.2300 | **0.0001** |
| Elevated plus maze | | | | | | | | |
| Time in open arms | 2.9980 | ***0.0862*** | 0.6664 | 0.4161 | 3.4900 | ***0.0644*** | 0.2026 | 0.6535 |
| Percent time in open arms | 3.2820 | ***0.0728*** | 1.0800 | 0.3010 | 3.3280 | ***0.0709*** | 0.2302 | 0.6324 |
| Percent entries in open arms | 0.0342 | 0.8537 | 0.3553 | 0.5524 | 0.2089 | 0.6486 | 0.0518 | 0.8294 |
| Three-chamber social interaction | | | | | | | | |
| Habituation ratio | 0.0216 | 0.8837 | 0.2151 | 0.6446 | 0.0015 | 0.9689 | 2.4580 | 0.1227 |
| Social preference ratio | 0.6262 | 0.4321 | 0.0077 | 0.9303 | 0.3275 | 0.5694 | 0.2348 | 0.6299 |
| Social memory ratio | 1.8400 | 0.1804 | 0.7987 | 0.3753 | 0.5268 | 0.4710 | 0.0610 | 0.8057 |
| Prepulse inhibition of acoustic startle | | | | | | | | |
| Baseline startle response | 0.3935 | 0.5318 | 0.0646 | 0.7999 | 1.0500 | 0.3078 | 3.9640 | **0.0485** |
| Percent average prepulse inhibition | 0.8726 | 0.3523 | 0.5551 | 0.4579 | 0.6383 | 0.4161 | 1.163 | 0.2833 |

**Notes**. Results from ANOVAs (treatment x lighting x sex) for behavioral parameters from poly IC- and saline mice, while also highlighting the treatment x sex interactions and main effects of sex analysis. ANOVA factors include sex (male, female), treatment (poly IC, saline exposure) and lighting (LD, LL). P-values in **bold** represent significance, and p-values in ***bold and italics*** represent trending significance.

**Supplementary Table 2. Sex differences in microglia of poly IC and saline-exposed mice under LD or LL.**

| | | Treatment x Sex x Lighting | | Treatment x Sex | | Lighting x Sex | | Main Effect Sex | |
|---|---|---|---|---|---|---|---|---|---|
| | | F | *p-value* | F | *p-value* | F | *p-value* | F | *p-value* |
| Dentate gyrus | | | | | | | | | |
| | Morphological index | 0.1327 | 0.7180 | 6.1890 | **0.0183** | 0.0158 | 0.9009 | 4.1630 | **0.0496** |
| | Cell body area | 0.7995 | 0.3778 | 0.0829 | 0.7752 | 0.3664 | 0.5492 | 10.8500 | **0.0024** |
| | Cell body circularity | 0.2634 | 0.6113 | 3.3340 | ***0.0772*** | 1.2580 | 0.2703 | 0.0518 | 0.8213 |
| | Density | 0.3323 | 0.5683 | 5.0250 | **0.0320** | 0.2974 | 0.5893 | 0.0343 | 0.8542 |
| | Nearest neighbor distance | 0.3788 | 0.5426 | 1.0760 | 0.3073 | 3.2630 | ***0.0803*** | 0.0439 | 0.8353 |
| | Spacing index | 0.4053 | 0.5289 | 1.7180 | 0.1993 | 3.4310 | ***0.0732*** | 0.0087 | 0.9263 |
| PFC | | | | | | | | | |
| | Morphological index | 0.3848 | 0.5394 | 0.6615 | 0.4220 | 2.7310 | 0.1082 | 6.4700 | **0.0160** |
| | Cell body area | 0.0093 | 0.9239 | 0.1080 | 0.7446 | 0.2653 | 0.6101 | 3.0760 | ***0.0890*** |
| | Cell body circularity | 0.0140 | 0.9065 | 0.1434 | 0.7074 | 0.2002 | 0.6576 | 1.0220 | 0.3196 |
| CA1 | | | | | | | | | |
| | Morphological index | 0.1226 | 0.7285 | 0.0115 | 0.9153 | 0.3753 | 0.5445 | 0.1210 | 0.7302 |
| | Cell body area | 0.0922 | 0.7634 | 0.2035 | 0.6549 | 3.9080 | ***0.0567*** | 18.7600 | **0.0001** |
| | Cell body circularity | 0.0480 | 0.8280 | 0.0269 | 0.8709 | 0.8782 | 0.3557 | 2.6610 | 0.1127 |
| | Density | 0.1363 | 0.7144 | 0.0100 | 0.9208 | 0.0020 | 0.9649 | 2.0220 | 0.1647 |
| | Nearest neighbor distance | 0.5015 | 0.4840 | 2.5400 | 0.1208 | 0.1212 | 0.7300 | 2.1790 | 0.1497 |

| | | | | | | | | |
|---|---|---|---|---|---|---|---|---|
| Spacing index | 1.932 | 0.1742 | 2.623 | 0.1152 | 0.155 | 0.6965 | 0.1436 | 0.7073 |

**Notes**. Results from ANOVAs (treatment x lighting x sex) for microglia parameters from poly IC- and saline mice, while also highlighting the treatment x sex interactions and main effects of sex analysis. ANOVA factors include sex (male, female), treatment (poly IC, saline exposure) and lighting (LD, LL). P-values in **bold** represent significance, and p-values in ***bold and italics*** represent trending significance.

**List of Supplementary Files**

**Supplementary File 1. Lists of differentially expressed genes (DEGs) in the dorsal hippocampus of male mice.** DEGs are listed in different tabs of the file, for the different comparisons between the four group (saline/LD, saline/LL, poly IC/LD, poly IC/LL).

**Supplementary File 2. Lists of differentially expressed genes (DEGs) in the dorsal hippocampus of female mice.** DEGs are listed in different tabs of the file, for the different comparisons between the four group (saline/LD, saline/LL, poly IC/LD, poly IC/LL).

**Supplementary File 3. List of genes in modules significantly associated with poly IC or LL exposure in males.** Genes are listed in different tabs of the file, for each of the 10 WGCNA modules associated with poly IC or LL exposure in males.

**Supplementary File 4. List of genes in modules significantly associated with poly IC exposure in females.** Genes are listed in different tabs of the file, for each of the 4 WGCNA modules associated with poly IC exposure in females (no module significantly associated with LL in females).

**Supplementary File 5. List of hub-bottleneck genes in the six WGCNA modules of Figure 9.**
